# Cooperative behavior in rat dyads reveals distinct social coordination strategies of increasing complexity

**DOI:** 10.1101/2025.09.24.678380

**Authors:** Ashutosh Shukla, Edward L. Rivera, John H. Bladon, Shantanu P. Jadhav

**Author notes:** Correspondence (S.P.J.). These authors contributed equally to the work (Ashutosh Shukla, Edward L. Rivera).

## Abstract

Cooperative behavior, the ability of individuals to coordinate their actions toward shared goals, is fundamental to survival and social success across species. However, the mechanisms supporting cooperation and how their disruption leads to social deficits in neurodevelopmental disorders such as autism spectrum disorder (ASD), remain poorly understood. To address these questions, we developed a novel cooperation task in paired spatial mazes under deterministic (100%) and probabilistic (50%) reward contingencies, utilizing dyads of littermate wild-type (WT) and *Fmr1* knockout (*Fmr1*) rats, a model of Fragile X syndrome. Both WT and *Fmr1* rat pairs exhibited dynamic turn-taking with mixed leader-follower behavior for cooperation; however, WT pairs achieved significantly greater cooperation success than *Fmr1* pairs. WT and *Fmr1* pairs both successfully utilized a simple follower-tracking-leader reactive strategy, dependent on partner-directed visual attention, resulting in slower asynchronous transitions, with *Fmr1* pairs more reliant on this strategy. WT rat pairs in addition exhibited a more efficient, flexible predictive cooperation strategy, dependent on coordination of optimal transition patterns between rat pairs and recent partner choice history, resulting in faster synchronous transitions. *Fmr1* rats were unable to show adaptation to the probabilistic reward condition by employing this more efficient social reciprocity strategy of higher complexity, leading to deficits in cooperative behavior. These findings reveal key behavioral strategies of varying complexity for cooperation, reveal the specific disruption of complex social reciprocity strategies in a rat ASD model, and provide a foundation for investigation of social interactions and underlying neural mechanisms in cooperative decision-making with relevance to neurodevelopmental disorders.

## INTRODUCTION

Cooperation is essential for survival and group cohesion in social species, including humans, enabling collective defense, resource sharing, and learning (Axelrod and Hamilton, 1981; Wilson, 2000; Krause and Ruxton, 2002; Nowak, 2006). These complex skills develop through observation and coordination within social groups, characterizing decision-making in natural environments (Giraldeau and Caraco, 2018). Under naturalistic settings, individuals constantly integrate information about their own goals with the actions of others in continuous reciprocal interactions, navigating interdependent and dynamic social contexts.

Successful cooperative interactions involve predicting partners’ actions, adjusting one’s own behavior in real time, and coordinating toward shared goals. These features make cooperation particularly informative for understanding the mechanisms of dynamic social decision-making. Moreover, impairments in social communication and flexible interaction with others are central characteristics of autism spectrum disorders (Boucher, 1977; Soderstrom, Rastam and Gillberg, 2002; Budimirovic *et al*., 2006; Salcedo-Arellano *et al*., 2020), especially, Fragile X syndrome, highlighting its value as a tractable framework for studying the biological and cognitive bases of social impairments.

Investigating behavioral principles and strategies for social cooperation can provide key insights into social cognition and inform intervention strategies. However, existing paradigms for studying cooperative behavior remain limited. Many rely on turn-based games or simplified laboratory tasks with fixed choice structures (Gardner *et al*., 1984; Avital, Aga-Mizrachi and Zubedat, 2016; Wood, Kim and Li, 2016; Delmas, Lew and Zanutto, 2019; Schweinfurth and Taborsky, 2020; Jiang *et al*., 2021; Shin and Ko, 2021; Zhang *et al*., 2023), which constrain opportunities for dynamic coordination, partner tracking, and reciprocal engagement. Developing ethologically grounded experimental designs that preserve the richness and flexibility of real-world social interactions is therefore critical for advancing our understanding of cooperative decision-making and its neural underpinnings.

Rats can be an ideal rodent model for these investigations due to their advanced socio-cognitive abilities and natural prosocial behaviors, including reciprocity, group foraging, and coordinated decision-making (Nagy *et al*., 2020, 2024; Schweinfurth, 2020). Rat foraging behavior in spatial contexts has been extensively studied since Tolman’s early experiments demonstrating cognitive maps (Tolman, 1948), and rats also tend to show more complex social play behavior such as hide-and-seek (Reinhold *et al*., 2019; Kaufmann, Brecht and Ishiyama, 2022). Importantly, several prior studies have attempted to probe cooperative behaviors in rats (Daniel, 1942; Rosenbaum and Epley, 1971; Schuster, 2002; Avital, Aga-Mizrachi and Zubedat, 2016), underscoring their capacity for socially motivated interactions. Rats’ advanced cognitive abilities, the rich repertoire of social behaviors and the availability of monogenic rat models with targeted gene knockouts such as *Fmr1* and *Nlgn3* (Hamilton *et al*., 2014; Till *et al*., 2015; Saxena *et al*., 2018; Asiminas *et al*., 2019; Klibaite *et al*., 2025), can provide a framework for investigating mechanisms underlying social coordination.

Here, we introduce a novel spatial cooperation task for rat dyads navigating two opposing W-mazes, in which animals must coordinate with a social partner to obtain food rewards by synchronizing visits to complementary maze reward locations, even under probabilistic reward conditions. This task takes advantage of rats’ natural spatial foraging behavior for rewards and enables intentional, real-time coordination. We show that efficient cooperation in this task can be accomplished by reciprocal coordination of choice strategies with peers, and through real-time monitoring using peer-directed attention which guide reactive actions. Further, wild-type (WT) littermate pairs employ a more efficient predictive coordination strategy to execute cooperation under probabilistic reward contingencies, while Fragile X knockout littermate rats (*Fmr1*) are deficient in this complex social reciprocity strategy. By quantifying coordination strategies, timing, and decision dynamics in both wild-type and *Fmr1* knockout rats, our findings provide a foundation for understanding the behavioral principles of cooperation, investigating the neural bases of cooperation, and how these processes may be disrupted in neurodevelopmental disorders.

## RESULTS

### A spatial cooperation task for social coordination in rat pairs

We developed a spatial cooperation task in which littermate rat pairs (dyads) from a Fragile-X knockout (*Fmr1)* rat colony must simultaneously choose matching reward wells from three options on complementary W-mazes separated by transparent barriers. Each rat navigates its own maze, but must learn to coordinate actions with its partner on the other maze to visit matching reward locations on complementary arms of the two mazes, and simultaneously nose-poke in the reward wells to receive automatically dispensed rewards (**Fig. 1A, 1B**, **see Methods**).

**Figure 1.**
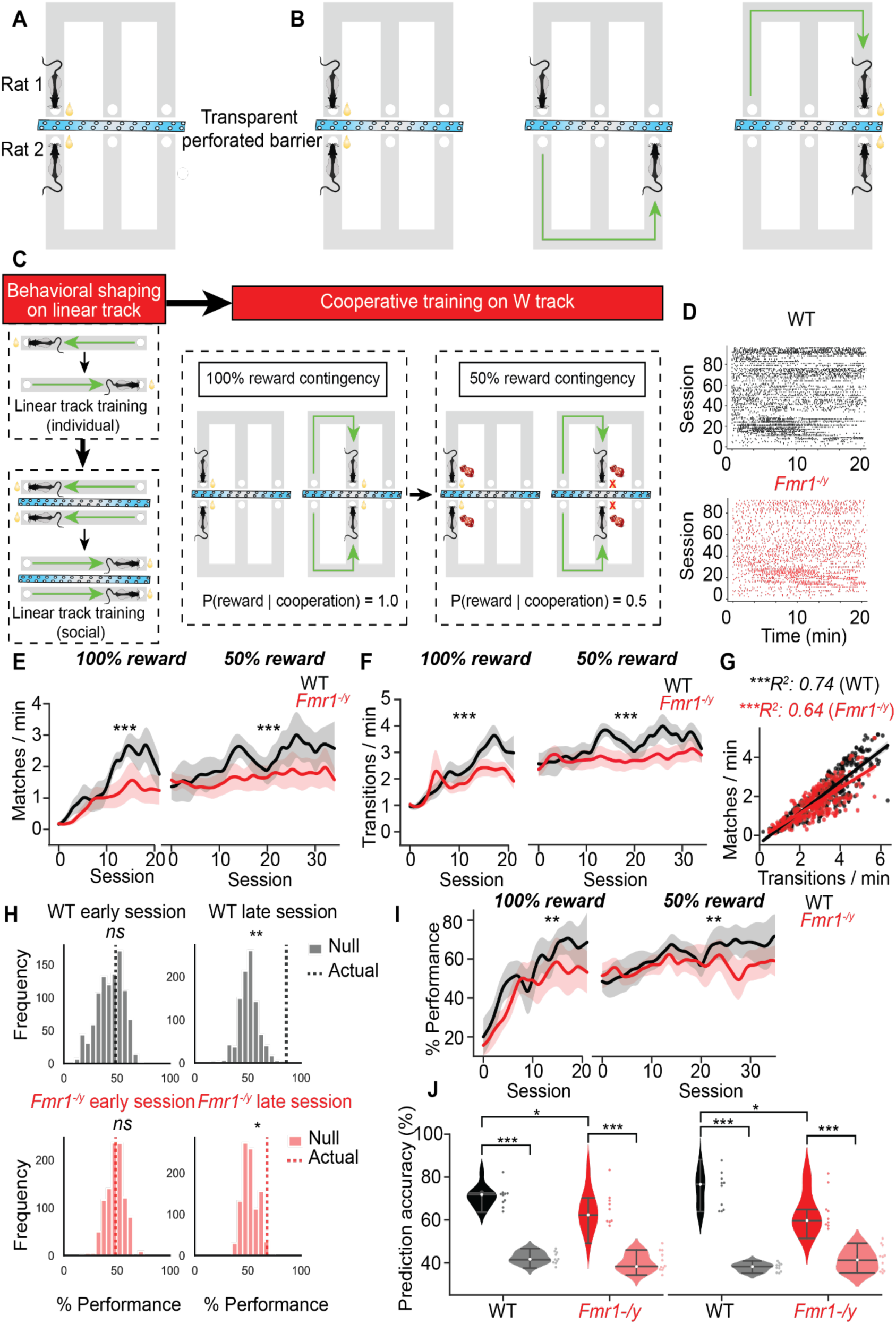
Cooperative learning task and coordination efficiency over training. A. Experimental setup. A pair of rats run on complementary W-mazes separated by a transparent perforated barrier to seek milk rewards. B. Example of a successful cooperative event. Both rats occupy complementary wells on their respective mazes (spatial cooperation) and poke simultaneously together in their respective wells (temporal cooperation) to collect milk reward (*left*). Following a successful match event, rat 1 moves to another well on the maze (*middle*). A successful cooperative event occurs if rat 2 chooses to go to the complementary reward well on its maze and poke simultaneously with rat 1 in the respective reward wells (*right*). C. Experimental design. *Left:* Rats were trained to run back and forth on a linear track alone (individual phase) and with another rat (social cooperation phase). Following proficiency in both the individual and social phases on the linear tracks, rats were trained on the social cooperative learning paradigm on 2 reward contingencies in succession: 100% reward contingency (where every cooperative event was rewarded), followed by 50% reward contingency (where cooperative events were rewarded probabilistically only 50% of the time). D. Raster plot showing match events for an example WT (*left*) and *Fmr1^-/y^* (*right*) rat pair. E. Match rates for WT and *Fmr1^-/y^* pairs for 100% (*left*) and 50% (*right*) reward contingencies across behavior sessions. *100%:* Two-way RM ANOVA (genotype: p < 0.05, session: p < 0.05, genotype X session: p > 0.05). *50%:* Two-way RM ANOVA (genotype: p < 0.05, session: p > 0.05, genotype X session: p > 0.05). * denote genotype-specific differences, *** p < 0.001. F. Transition rates for WT and *Fmr1^-/y^* rats under 100% (*left*) and 50% (*right*) reward contingencies. *100%:* Two-way RM ANOVA (genotype: p < 0.001, session: p < 0.001, genotype X session: p > 0.05). *50%:* Two-way RM ANOVA (genotype: p < 0.001, session: p > 0.05, genotype X session: p > 0.05). *** p < 0.001. G. Correlation between cooperative match events and transition rates. Two-sided t-test. *** p < 0.001. H. *Top:* WT pair: Observed vs. shuffled coordination performance across training. Null distributions of coordination performance for a representative WT pair during an early session (100% contingency, *left*) and late (50% contingency, *right*) session are shown. Vertical dashed lines indicate observed values. Null distributions were generated from 1,000 circular time-shift permutations of one animal’s behavioral sequence, preserving individual choice patterns while disrupting inter-animal timing. Observed performance did not exceed chance in early sessions (*ns*) but significantly surpassed the null distribution in late sessions (p < 0.01), indicating learned coordination. *Bottom: Fmr1*^−/y^ pair: Observed vs. permuted coordination performance across training. Same analysis for a representative *Fmr1*^−/y^ pair. Observed coordination did not differ from chance in early sessions (*ns*) but showed a modest, yet significant increase above the null distribution in late sessions (p < 0.05), suggesting limited improvement in cooperative timing with training. I. Performance metrics for WT and *Fmr1^-/y^* pairs in 100% (*left*) and 50% (*right*) reward contingencies shows significant impairment in *Fmr1^-/y^* pairs compared to WT pairs. *100%:* Two-way RM ANOVA (genotype: p < 0.01, session: p < 0.001, genotype X session: p > 0.05). *50%:* Two-way RM ANOVA (genotype: p < 0.01, session: p > 0.05, genotype X session: p > 0.05). ** denote genotype-specific differences, ** p < 0.01. J. Prediction accuracy of rats’ next choice given partner’s current choice in 100% (*left*) and 50% (*right*) reward contingencies (Mann-Whitney U-test, * p < 0.05, *** p < 0.001).

In a behavioral shaping step, rat pairs were trained on parallel linear tracks separated by a transparent, perforated barrier and rewarded for simultaneous nose-pokes in matching reward wells, facilitating the transition from individual to coordinated actions (**Fig. 1C**). Rat pairs were then introduced to the spatial cooperative task on paired W-mazes, first with a deterministic 100% reward contingency for each *“match”/coordinated event* of nose-poking on matched reward wells, resulting in both rats receiving rewards in their respective wells for every successful match event. After training on several sessions with 100% rewards over multiple days, the task was then switched to a 50% probabilistic reward contingency, in which the rats were only rewarded with a random 50% chance for each successful match event (**Fig. 1C**, examples of successful and unsuccessful cooperation shown in **Movie 1** and **Movie 2** respectively). Rats performed at least two behavioral sessions per day interleaved with a rest/ sleep epoch, and each individual rat is run on both mazes within a day (switching assigned maze after each session). To assess behavioral flexibility and promote optimal exploration of all three maze arms, we trained rat pairs on the 50% probabilistic reward contingency. For the deterministic 100% contingency, each match event leads to reward, so rat pairs can choose run back and forth between any two reward well arms of their maze in a coordinated manner, similar to the linear maze. In contrast, in the 50% probabilistic condition, after each successful rewarded “match” event on a given maze arm, only one of the remaining two arms was randomly chosen as the reward-arm for the next transition. Both animals therefore must together explore the remaining two possible options until they get rewarded, encouraging optimal exploration of all three arms of the mazes. There is 50% probability of match events being rewarded, and animals must continue coordinating actions (spatial trajectories), even if they do not get rewarded after a match event, requiring cooperative behavior irrespective of reward success. Coordination of well-visits on opposing arms therefore becomes a primary goal of the task, which must be learnt by rat pairs together.

Adult male littermate rat pairs (dyads)–WT-WT (WT pairs), *Fmr1-Fmr1* (*Fmr1* pairs), and WT-*Fmr1* (mixed pairs), with *N* = 5 pairs per group–were trained on this cooperative task. Cooperative behavior learning performance over sessions for WT and *Fmr1* pairs for both the 100% and 50% contingencies are shown in **Fig. 1D**. Both WT and *Fmr1* rat pairs exhibited performance indistinguishable from chance during early training on both reward contingency tasks, however, by late training, their performance improved significantly, exceeding chance levels (**Fig. 1H, 1I**). Importantly, WT pairs demonstrated significantly higher coordination rates than both *Fmr1* pairs (**Fig. 1E, 1I**) and mixed pairs (**Fig. 1–Supplementary 1A**) under both 100% and 50% reward contingencies. *Fmr1* and mixed pairs were generally slower in the task, as reflected by a reduced number of task-related transitions (**Fig. 1F, G, Fig. 1−Supplementary 1D**), but even accounting for this difference, their learning and performance in cooperative behavior was impaired (**Fig. 1**, **Fig. 1 Supplementary 1−B**). Consistent with enhanced learning and performance, the current location of a peer provided sufficient information to reliably predict a rat’s subsequent response (**Fig. 1J**, **Fig. 1 Supplementary 1−C**). This predictive relationship exceeded chance levels across all rat pairs. However, prediction accuracy was significantly higher in WT pairs compared to both Fmr1 and mixed-genotype pairs, highlighting a genotype-dependent modulation of social predictive cues. Taken together, these results demonstrate that while all rat pairs were capable of learning to coordinate, WT pairs exhibited superior cooperative performance, faster transitions in trajectories between reward wells, and enhanced sensitivity to peer cues. These findings suggest that *Fmr1* rats exhibit specific impairments in the acquisition and use of social information critical for effective cooperation.

### Cooperation deficits in *Fmr1* rats are not due to sensory or learning deficits

To further isolate the factors that contribute to successful coordination in WT rats, and the impairment of such coordination in *Fmr1* rats, we conducted a series of targeted control experiments. First, to determine the contribution of social vision, we replaced the transparent, perforated divider with an opaque barrier (No Vision condition / NV) that prevented exchange of visual information between rats (**Fig. 2A**), testing whether successful cooperation could be achieved through either random transitions between maze arms or through use of non-visual strategies such as rhythmic coordinated transitions between maze arms. These sessions were interspersed with the standard 50% reward condition sessions, during which the rats could see each other. Compared to the 50% reward condition, rats showed a significant reduction in performance in the NV sessions (**Fig. 2B, C**), demonstrating that visual access to the partner, and possibly peer-directed attention, is necessary for successful joint coordination. Importantly, all experiments were conducted in the presence of continuous auditory white noise to mask potential auditory communication, thereby ruling out ultrasonic vocalizations as a primary modality for coordination. Second, to assess whether the lower cooperation performance could be due to differences in associative learning (associating partner presence with reward), since memory deficits have been reported in *Fmr1* rats (Asiminas *et al*., 2019; Fernandes *et al*., 2021), we tested whether WT and *Fmr1* rats could form similar associations in a non-social context by replacing the social partner with a localized visual cue presented at one of the reward wells. (**Fig. 2D**). As in the main task, this control was conducted under both 100% and 50% reward contingencies. Both WT and *Fmr1* rats learnt to approach the reward-associated arm based on the visual cue (**Fig. 2E**), demonstrating intact visual cue-reward associative learning. *Fmr1* rats did exhibit slower learning rates (**Fig. 2E**) and delayed approach behavior compared to WT rats (**Fig. 2F**), reflecting differences in task performance speed, rather than deficits in learning or sensory processing, that likely contributes to their slower performance in the cooperation task. Together, these results underscore the crucial role of visual cues in supporting successful coordination in WT, *Fmr1*, and mixed pairs, while demonstrating that *Fmr1* rats retain visual processing and visual- reward association capabilities, indicating that deficits in these factors cannot solely account for their impaired performance in the cooperative foraging task.

**Figure 2.**
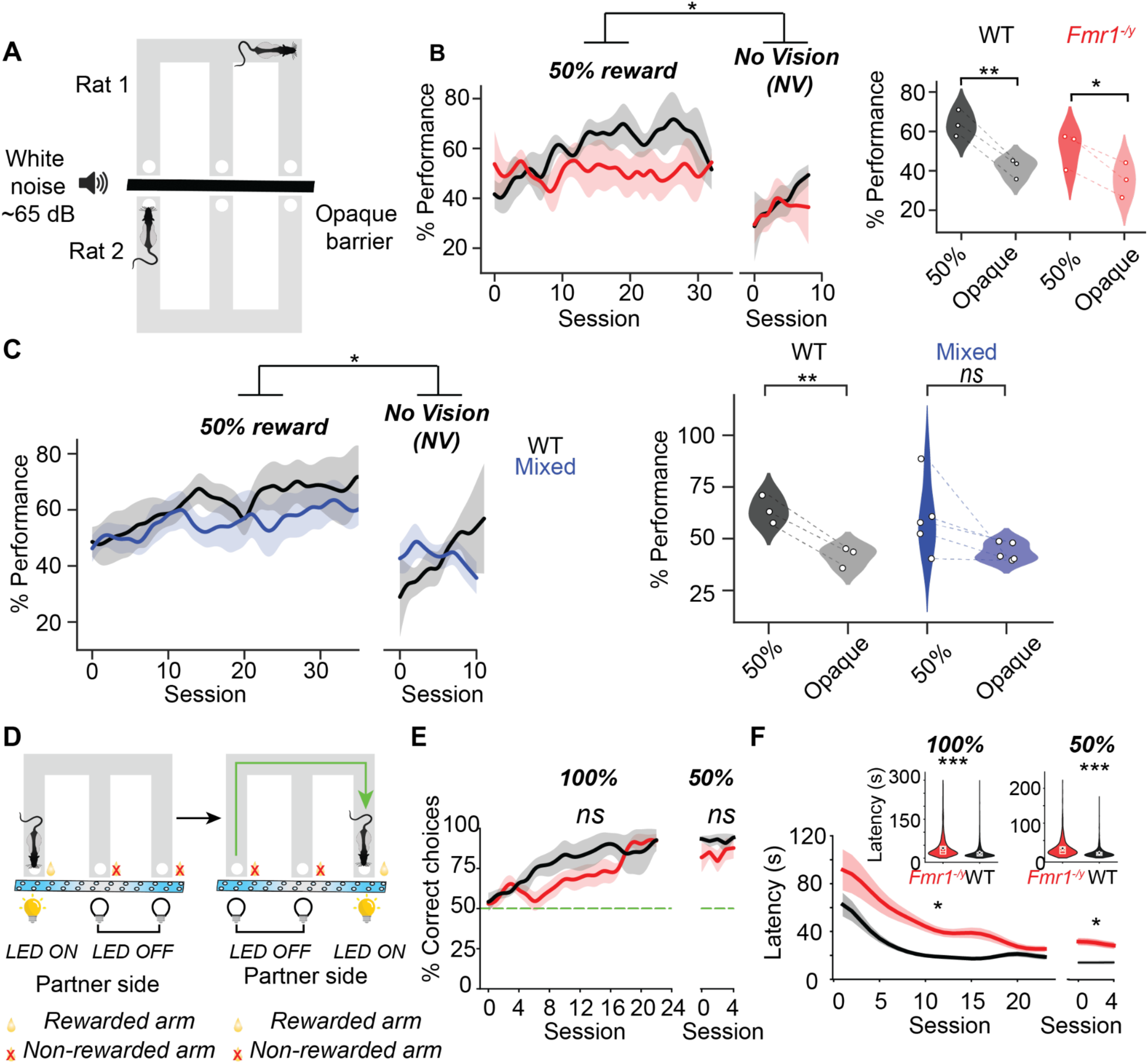
Behavioral controls. A. Schematic showing No Vision (NV) control sessions. Rats in a pair ran on their respective mazes with an opaque barrier between mazes obscuring sight. B. Session wise comparison of performance of rat pairs between 50% reward (WT: N = 3 pairs; *Fmr1^-/y^*: N = 3 pairs) and no vision conditions. Two-way RM ANOVA (genotype: p > 0.05, condition: p < 0.05, genotype X condition: p > 0.05). *Right:* Comparison of mean performance of rat pairs between transparent (last 9 sessions, N = 3 pairs) and opaque conditions (N = 3 pairs). Paired t-test,* p < 0.05, ** p < 0.01. C. Session wise comparison of performance of rat pairs between 50% reward (WT: N = 3 pairs; Mixed: N = 5 pairs) and no vision conditions. Two-way RM ANOVA (genotype: p > 0.05, condition: p < 0.05, genotype X condition: p > 0.05). *Right:* Comparison of mean performance of rat pairs between transparent) and opaque conditions. Paired t-test. *ns*: not significant. ** p < 0.01. D. Schematic for visual cue association task. Rats were trained on a W-maze to approach an LED-cued arm on the opposite maze to seek milk rewards on the corresponding, apposing reward well on it’s own maze. E. Performance comparison between WT and *Fmr1^-/y^* rats on the visual cue-association task under 100% (*left*) and 50% (*right*) reward contingencies. *100%:* Mixed effects model (genotype: p > 0.05, session: p > 0.05, genotype X session: p > 0.05). *50%:* Mixed effects model (genotype: p > 0.05, session: p < 0.001, genotype X session: p > 0.05). *ns*: not significant. F. *Top:* WT rats respond to the visual LED cue with lower latency relative to *Fmr1^-/y^* rats under both 100% (*left*) and 50% (*right*) reward contingencies. 100%: *Fmr1^-/y^* : *n =* 2,371, *mean* = 42.2 s; WT: *n =* 4,554, *mean* = 22.0 s; *p* = 6.9×10⁻²²⁷, Mann-Whitney U test). 50%: *Fmr1^-/y^*: *n* = 862, *mean* = 28.4 s; WT: *n = 1*,359, *mean* = 14.0 s; *p* = 1.3×10⁻⁸², Mann-Whitney U test). *Bottom:* Latency to respond to visual cue for sessions under 100% (*left*) and 50% (*right*) reward contingencies. *100%:* Mixed effects model (genotype: p < 0.05, session: p < 0.001, genotype X session: p < 0.01). *50%:* Mixed effects model (genotype: p < 0.05, session: p < 0.01, genotype X session: p > 0.05).

### Development of structured spatial and temporal coordination strategies supports cooperative behavior

We next examined whether rats developed structured strategies that supported efficient joint exploration, a critical component of coordinated behavior. Beyond moment-to-moment monitoring and reactive actions based on partner location, successful cooperation may require higher-order planning across extended sequences. To address this, we analyzed how pairs coordinated both the spatial structure of their exploration and the temporal alignment of their actions.

We first compared exploration patterns across reward contingencies with differing task demands. Under the 100% reward condition, coordinated alternation between any two arms was sufficient to maximize reward, whereas the 50% probabilistic condition required systematic sampling of all three arms for optimal exploitation. Reward was maximized when rats alternated through one of two optimal arm sequences (maze arm transitions 1–2–3 or 1–3–2; **Fig. 3A**). In line with these differing requirements, transition probabilities between maze arms showed distinct patterns across contingencies (**Fig. 3B**). Parsing each session’s transitions into non-overlapping triplets revealed that rats in the 100% condition predominantly alternated between two adjacent arms, while those in the 50% condition engaged in comprehensive exploration of all three arms through either the 1–2–3 or 1–3–2 sequences, especially for WT pairs (**Fig. 3B, 3C**). Crucially, WT pairs exhibited greater coordination in executing optimal triplet sequences than *Fmr1* pairs (**Fig. 3C, 3E**). WT rats also flexibly alternated between the two optimal sequences within and across sessions and adapted strategies according to task demands (**Fig. 3C**). Further, the entropy of their choice sequences declined over training, reflecting increasing regularity and efficiency (**Fig. 3D**).

**Figure 3.**
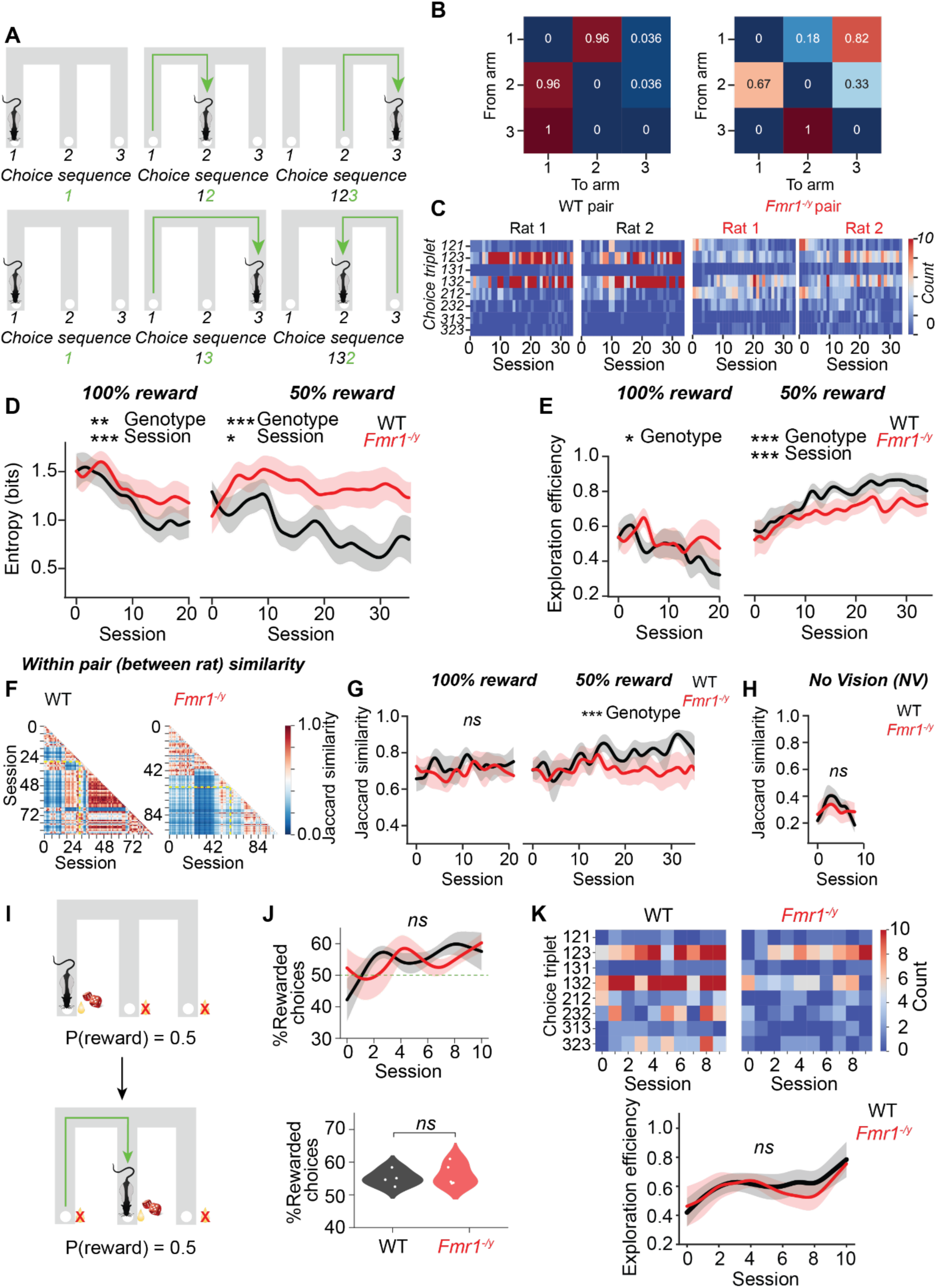
Emergence of optimal spatial coordination strategies. A. Two example spatial trajectory choice sequences (*top* and *bottom*) that lead to optimal sampling of available reward space. B. Well-to-well transition probabilities from an example WT session under 100% (*left*) and 50% (*right*) reward contingencies. Note the change in transition probabilities across the two reward contingencies. C. Counts of unique choice triplets in 50% reward contingency sessions for rats from a representative WT pair and a representative *Fmr1^-/y^* pair. D. Session wise comparison of entropy of choice sequences of WT and *Fmr1^-/y^* rats during 100% (*left*) and 50% (*right*) reward contingencies, showing significantly higher decrease in entropy with learning for WT rats. N = 10 rats each for WT and *Fmr1^-/y^*; *100%:* Two-way RM ANOVA (genotype: p < 0.01, session: p < 0.001, genotype X session: p > 0.05). *50%:* Two-way RM ANOVA (genotype: p < 0.001, session: p < 0.05, genotype X session: p > 0.05). ** p < 0.01, *** p < 0.001. E. Exploration efficiencies of WT and *Fmr1^-/y^* rats during 100% (*left*) and 50% (*right*) contingency sessions, showing significantly higher exploration efficiency for WT rats in the 50% condition. *100%:* Two-way RM ANOVA (genotype: p < 0.05, session: p > 0.05, genotype X session: p > 0.05). *50%:* Two-way RM ANOVA (genotype: p < 0.001, session: p < 0.001, genotype X session: p > 0.05). F. Heatmaps of Jaccard similarity between choice sequences, shown between partners within and across sessions for an example WT pair and an example *Fmr1^-/y^* pair. Horizontal and vertical dashed yellow lines indicate the transition from the 100% to 50% reward contingency. G. Session wise comparison of Jaccard similarity of choice sequences rats within WT and *Fmr1^-/y^* pairs during 100% (*left*) and 50% (*right*) reward contingencies, showing significantly higher decrease in entropy with learning for WT rats. N = 5 pairs each for WT and *Fmr1^-/y^*; *100%:* Two-way RM ANOVA (genotype: p > 0.05, session: p > 0.05, genotype X session: p > 0.05). *50%:* Two-way RM ANOVA (genotype: p < 0.001, session: p > 0.05, genotype X session: p > 0.05). *ns*: not significant. *** p < 0.001. H. Session wise comparison of Jaccard similarity of choice sequences rats within WT and *Fmr1^-/y^*pairs (N = 3 pairs per genotype) during no vision (NV) control conditions, showing no difference between genotypes. However, the reduction in Jaccard similarity relative to that shown in **G** is noteworthy. *ns*: not significant. I. Schematic for agent simulation experiments. Rats learned to transition between maze arms to receive rewards with reward contingency pre-determined by a computer program, similar to the 50% reward contingency in the cooperative foraging task. J. *Top:* Session wise performance of WT and *Fmr1^-/y^*rats in the agent simulation experiments were similar to each other. N = 4, 5 rats for WT and *Fmr1^-/y^* respectively; Two-way RM ANOVA (genotype: p > 0.05, session: p > 0.05, genotype X session: p > 0.05). *Bottom:* Mean performance was same across genotypes (Mann-Whitney U test, p > 0.05). *ns*: not significant. K. *Top:* Heatmap of unique choice triplet counts for an example WT (*left*) and *Fmr1^-/y^* (*right*) rat. *Bottom:* Exploration efficiency of WT and *Fmr1^-/y^*rats were similar (Two-way RM ANOVA, genotype: p > 0.05, session: p > 0.05, genotype X session: p > 0.05). *ns*: not significant.

Building on the finding that WT rats developed more structured and efficient individual exploration than their *Fmr1* counterparts, we next assessed coordination between partners within rat pairs. Under the 100% reward condition, Jaccard similarity (see **Methods**) between the choice sequences of paired rats was high for both genotypes (**Fig. 3F, G**). In contrast, under the 50% probabilistic condition, WT pairs exhibited significantly higher similarity in their choice sequences compared with *Fmr1* pairs (**Fig. 3F, G**), indicating that only WT rats converged on shared, structured strategies when task demands were elevated. Importantly, during the no-vision condition, where direct partner monitoring was eliminated, similarity decreased comparably across genotypes, suggesting that the enhanced coordination observed under normal social conditions was specifically driven by partner-based interactions (**Fig. 3H**).

To test whether these differences reflect coordination per se rather than baseline differences in exploration efficiency or cognitive ability, we introduced a control task in which rats coordinated with a virtual agent under a 50% reward contingency (**Fig. 3I**). In this task, both WT and *Fmr1* rats achieved similar levels of task performance with comparable learning rates (**Fig. 3J**). As in the 50% reward condition of the original social task, optimal outcomes required systematic alternation through the maze arms using either of the two triplet sequences (1-2-3 or 1-3-2). Notably, both genotypes independently demonstrated the capacity to adopt such structured transition strategies when interacting with the virtual agent (**Fig. 3K**), indicating that efficient and temporally extended exploration strategies were not impaired in *Fmr1* rats when peer coordination demands were not required. This suggests that WT rats exhibit flexibility in adapting their exploration strategy, likely through an internal model of partner strategy that can be updated as required using peer-monitoring , a process that is impaired in *Fmr1* rats.

Having established that WT and *Fmr1* rats differed in the spatial organization of their joint exploration strategy, we next asked whether they also differed in the temporal precision of their coordination. We first quantified the temporal dynamics of rats’ behavior based on their arrival and departure times at reward wells (**Fig. 4A, 4B**). To examine how these behavioral dynamics evolved over training, we quantified three temporal parameters: mean arrival lag between rats during match events (**Fig. 4C**), mean departure lag from wells following matches (**Fig. 4D**), and mean dwell time at wells during both match and non-match events (**Fig. 4E**). We observed that, under the 100% reward contingency, the mean arrival lag during match events progressively decreased in WT pairs, whereas it remained consistently elevated in *Fmr1* pairs (**Fig. 4C**). This difference persisted under the 50% reward contingency, with WT pairs exhibiting significantly shorter arrival lags than *Fmr1* pairs (**Fig. 4C**). A similar pattern emerged for mean departure lag following match events, with WT rats departing more promptly than *Fmr1* rats (**Fig. 4D**). Additionally, *Fmr1* rats exhibited longer dwell times at reward wells compared to WT rats (**Fig. 4E**). To further probe whether these differences in arrival, departure, and dwell time dynamics reflected broader patterns of temporal coordination, we next examined the fine-grained synchrony of transitions using cross-correlation analyses of arrival times of rats in a pair at reward wells (**Fig. 4F, 4G**). Cross-correlation of arrival times at reward wells revealed significant peaks at multiple lags for both groups, exceeding shuffled controls and indicating that pairs learned to align their transitions within characteristic temporal windows. Importantly, only WT rats showed prominent peaks near zero lag, reflecting near-synchronous transitions consistent with a gaze-independent strategy that emerged with training, indicating that rats in a pair arrive at reward wells in close temporal proximity, moving in tandem in synchronous movements/ trajectories. *Fmr1* rats lacked this short-lag synchrony, indicating a reliance on asynchronous movements and possibly a gaze dependent strategy involving a follower tracking a leader’s position followed by a reactive action to match positions. Correspondingly, WT rats exhibited shorter lags at peak correlation than *Fmr1* rats, especially under the 50% condition (**Fig. 4I**), and achieved higher peak correlation values across sessions (**Fig. 4J**).

**Figure 4.**
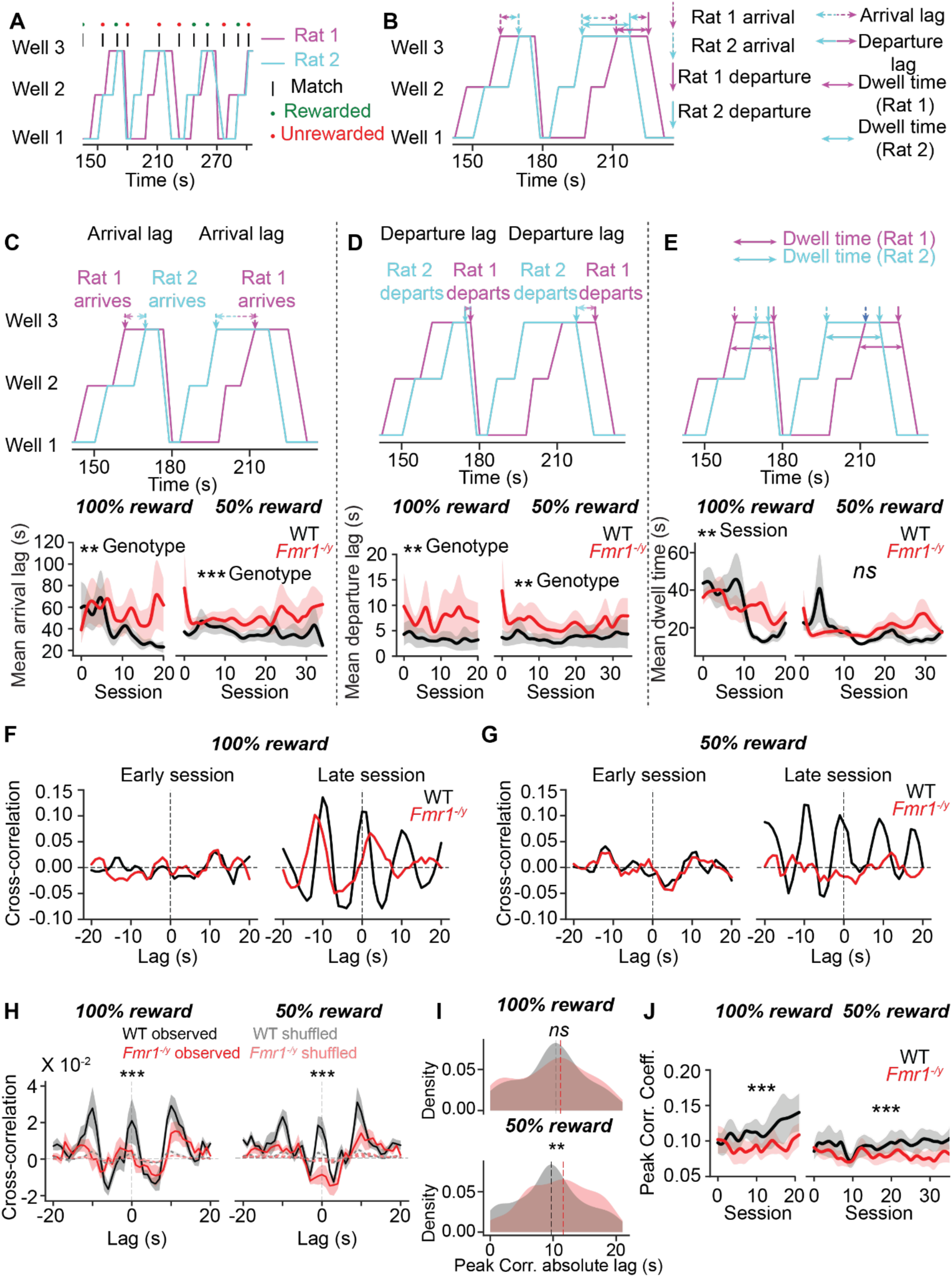
Emergence of temporal coordination and timescales of behavioral organization. A. A snippet of rat visit sequences from an example session showing well visit sequences for a rat pair. B. Zoomed in from panel A defining behavioral metrics that describe temporal dynamics of cooperative behavior. C. *Top: Left:* Snippet showing arrival lag between rats for a cooperative event. *Bottom: Left:* Mean arrival lag between rats in WT and *Fmr^1-/y^* pairs for 100% (*left*) and 50% reward contingencies (*right*). N = 5 pairs in each group; Two-way RM ANOVA (genotype: p < 0.01, session: p > 0.05, genotype X session: p > 0.05). *50%:* Two-way RM ANOVA (genotype: p < 0.001, session: p > 0.05, genotype X session: p > 0.05). *** p < 0.001. D. *Top: Middle:* Snippet showing departure lag between rats after a cooperative event. *Bottom: Middle:* Mean departure lag between rats in WT and *Fmr^1-/y^* pairs for 100% (*left*) and 50% reward contingencies (*right*). N = 5 pairs each; Two-way RM ANOVA (genotype: p < 0.05, session: p > 0.05, genotype X session: p > 0.05). *50%:* Two-way RM ANOVA (genotype: p < 0.01, session: p > 0.05, genotype X session: p > 0.05). ** p < 0.01. E. *Top: Right:* Snippet showing dwell time at a reward well for rats’ every visit. *Bottom: Right:* Mean dwell time at reward wells for WT and *Fmr^1-/y^*rats for 100% (*left*) and 50% reward contingencies (*right*). N = 10 rats in each group; Two-way RM ANOVA (genotype: p > 0.05, session: p < 0.01, genotype X session: p > 0.05). *50%:* Two-way RM ANOVA (genotype: p > 0.05, session: p > 0.05, genotype X session: p > 0.05). *ns*: not significant. ** p < 0.01. F. Example cross-correlation (smoothened) between well arrival times of an example WT and *Fmr1^-/y^*rat pair for a representative early (*left*) and late session (*right*) under 100% reward condition. G. Example Cross-correlation between well arrival times of an example WT and *Fmr1^-/y^* rat pair for a representative early (*left*) and late session (*right*) under 50% reward condition. Note the lack of systematic cross-correlation peaks for *Fmr1^-/y^* rat pairs in the 50% condition even after training. H. Cross-correlation coefficient values between well arrival times across 100% (*left*) and 50% (*right*) reward conditions for WT and *Fmr1^-/y^* pairs. These values were quantified for each pair session wise and averaged across pairs and sessions for both observed and shuffled data. *100%:* Linear mixed-effects model (genotype: p < 0.001, lag: p < 0.001, genotype X lag: p < 0.001). *50%:* Linear mixed-effects model (genotype: p < 0.001, lag: p < 0.001, genotype X lag: p < 0.001). *** p < 0.001. I. Distribution (kernel density estimates) of absolute lag values at peak cross-correlation for all sessions across 100% (*top*) and 50% (*bottom*) reward conditions for WT *Fmr1^-/y^* rat pairs. KS test (*100%:* p > 0.05, *50%:* p < 0.01). *ns*: not significant. ** p < 0.01. J. Session wise peak correlation coefficients for WT and *Fmr1^-/y^* rat pairs, showing greater temporal coordination for WT rat pairs. *100%:* Linear mixed-effects model (genotype: p < 0.001, session: p > 0.05, genotype X session: p > 0.05). *50%:* Linear mixed-effects model (genotype: p < 0.001, session: p > 0.05, genotype X session: p > 0.05). *** p < 0.001.

Together, these findings demonstrate that WT rats not only structured their exploration in space but also synchronized it in time, enabling efficient joint foraging. *Fmr1* rats showed deficits in both domains. They failed to organize exploration into structured sequences and were unable to coordinate transitions with short temporal lags. Overall, their behavior was organized at longer timescales relative to their WT counterparts. Further, the differences in structured exploration were exclusive to the social context of the cooperative foraging task and not to other non-social contexts as seen in the control experiments. These impairments highlight a specific vulnerability in the social decision-making mechanisms required for cooperative behavior.

### Dynamic turn-taking in the initiation of coordinated responses

Complementary leader–follower roles are amongst the most common strategies for organizing coordinated group behavior in social animals (King, Johnson and Vugt, 2009; Massen *et al*., 2010; Gachomba *et al*., 2022; Cheng *et al*., 2025; Meisner *et al*., 2025). Clear peaks at both negative and positive lags in the cross-correlation of arrival times prompted us to examine these dynamics more directly, focusing on sequences of coordinated match events across trials **(Fig. 5)**. For each match event, the rat that arrived first at the reward well was designated as the “Leader” and the peer as “Follower” for that event, with similar leader-follower designations for subsequent match events (**Fig. 5A, B**). The probability of maintaining the leader role across consecutive match events remained near chance levels for both WT and *Fmr1* rats over the course of learning (**Fig. 5C**), indicating flexible and dynamic turn-taking (example in **Movie 3**). On average, both genotypes led approximately half of the match events under both 100% and 50% reward contingencies. However, under the 50% reward condition, a small but significant genotype-specific difference emerged, with WT and *Fmr1* rats differing significantly in their likelihood of assuming the leader role at the start of training (**Fig. 5D**). However, this genotype-specific difference disappeared as training progressed. Furthermore, the likelihood of a rat assuming the leader role on a given trial was modulated by its leadership status over the preceding trials, indicating that leadership assignments fluctuate dynamically over time rather than remaining fixed (**Fig. 5−Supplementary 1C, D**). These findings demonstrate dynamic turn-taking behavior in WT and *Fmr1* rats in assuming leadership roles.

**Figure 5.**
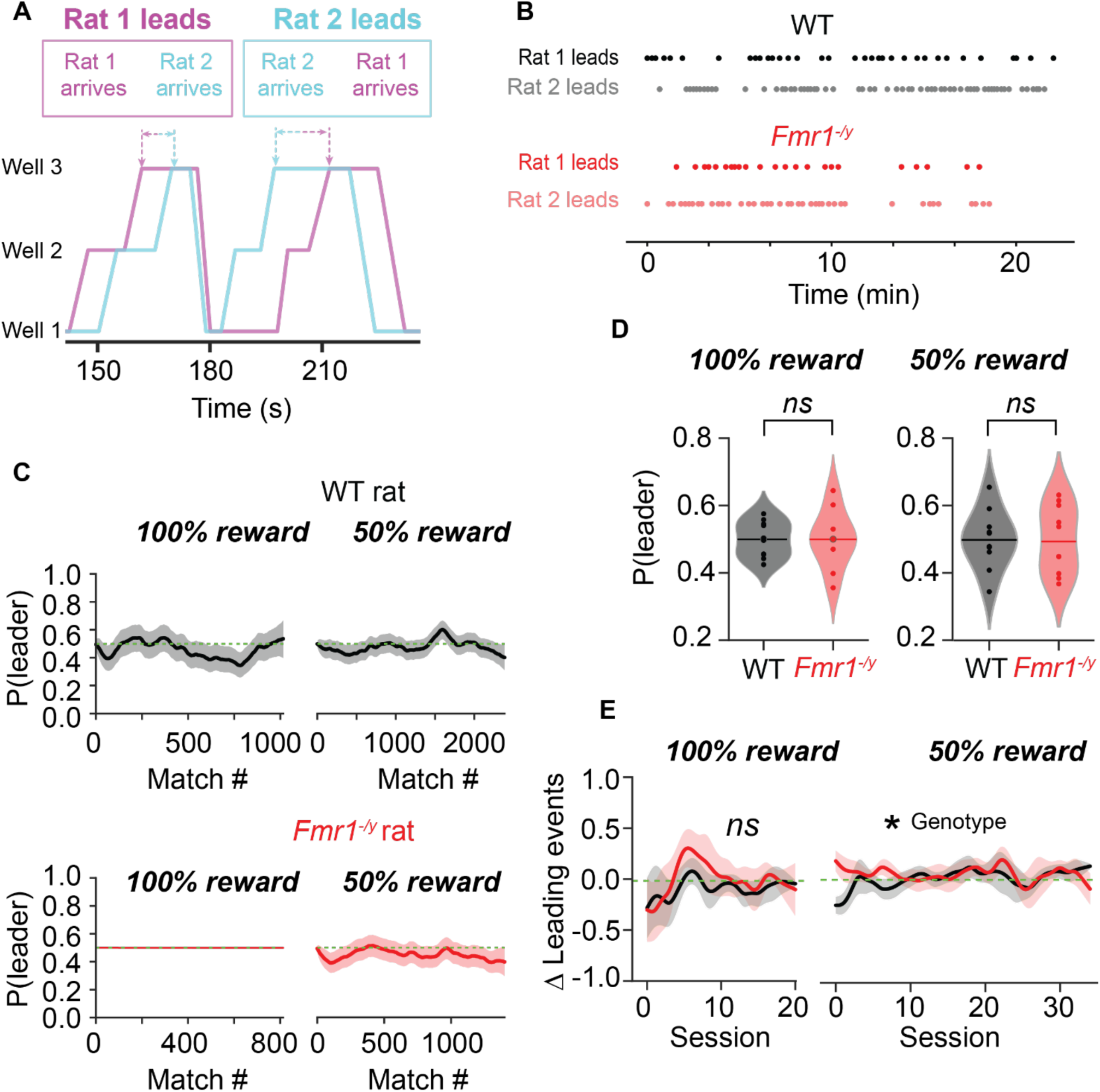
Dynamic leader-follower relationship during cooperative behavior. A. Snippet from an example session showing leader-follower relationship between rats in a pair for exemplar match events. B. Raster plot of leader assignments from an example session showing dynamic leader-follower relationships between rats of a WT pair (*top*) and a *Fmr1^-/y^* pair (*bottom*). C. Probability of assuming leader role for an example WT (*top*) and *Fmr1-/y* rat (*bottom*) within respective pairs as predicted by a state-space model under 100% (*left*) and 50% reward conditions (*right*). Green dashed horizontal line denotes probability of 0.5. D. Mean probability of assuming leader role (state space model) under 100% (*left*) and 50% (*right*) reward contingencies (Mann-Whitney U test, p > 0.05). *ns*: not significant. E. Difference in proportion of leading events between rats in a pair in 100% (*left*) and 50% reward contingencies (*right*). *100%:* Two-way RM ANOVA (genotype: p < 0.05, session: p > 0.05, genotype X session: p > 0.05). *50%:* Two-way RM ANOVA (genotype: p < 0.05, session: p > 0.05, genotype X session: p > 0.05). *ns*: not significant. * p < 0.05.

### Differential reliance on gaze-dependent strategies for successful coordination

Peer-directed social attention is a well-established hallmark of cooperative behavior across species, including primates and marmosets (Emery *et al*., 1997; Nahm *et al*., 1997; Franch *et al*., 2024; Meisner *et al*., 2025). Performance deficits in the no-vision (NV) condition of the cooperative foraging task indicated that vision-based social attention was essential for coordinating foraging across WT and *Fmr1* rat pairs. We hypothesized that rats strategically deploy social gaze (quantified using peer-directed head orientation) to gather task-relevant information and facilitate coordination. To examine this, we adopted a multipronged approach combining computational modeling and data-driven analyses. First, using a multi-agent reinforcement learning (MARL) framework (Tan, 1993; Du *et al*., 2023; Tsutsui *et al*., 2024), we evaluated the contribution of social attention to cooperative foraging. In these simulations, agent pairs foraged between three reward patches (analogous to the three reward wells in the maze) under three distinct policies: *partner-aware* (agents received partner-related information alongside environmental and self-related information), *partner-unaware* (agents received only environmental and self-related information), and *random* (agents received partner-related information alongside environmental and self-related information but made random choices) (**Fig. 6A**). Agents in a pair were required to coordinate their choices alike those in the cooperative foraging task, with both receiving a discounted reward on successful coordination. Like rat pairs, agent pairs started their training under 100% reward condition and after 100 training episodes switched to 50% reward condition for another 100 episodes. Partner-aware agent pairs achieved markedly higher performance than both partner-unaware and random agent pairs (**Fig. 6B**), underscoring the critical role of social attention in cooperative foraging.

**Figure 6.**
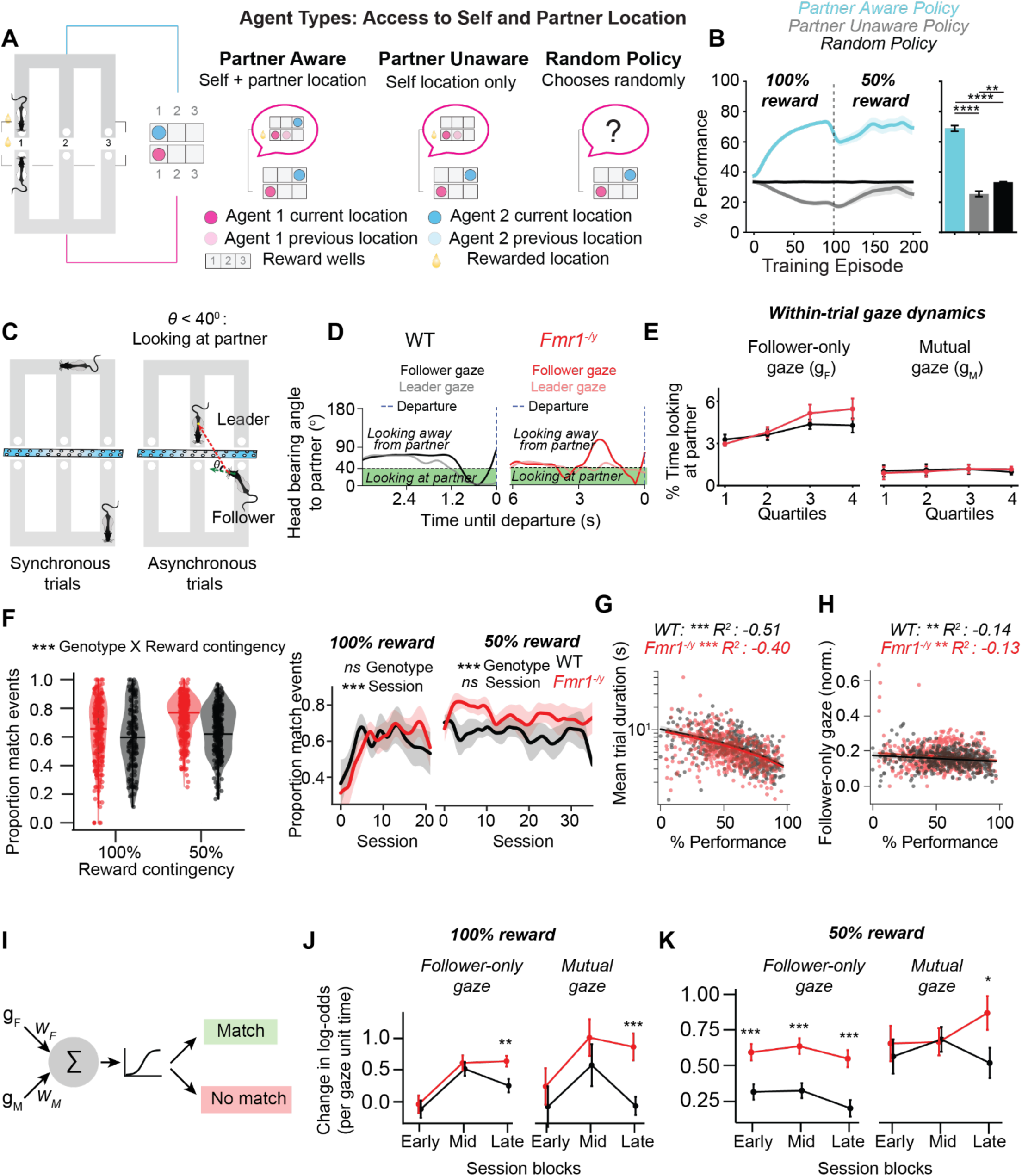
Role of partner-directed attention in successful cooperative behavior. A. Schematic of the spatial cooperation task (left) and corresponding cooperative multi-agent reinforcement learning (RL) framework (right). Each agent navigates its own discrete 3-location grid. B. Agent types differ in available task-relevant information. Partner-aware agents observe both their own and their partner’s current location. Partner-unaware agents observe only self-related cues. Random agents receive no observations and select among the three locations at random. Partner awareness enhances cooperative learning. Learning curves show cooperation efficiency (% of optimal matches) over 200 training episodes for Partner-Aware (cyan), Partner-Unaware (gray), and Random (black) agents. A reward contingency switch at episode 100 (dashed line) reduces the reward probability from 100% to 50%, similar to the spatial coordination task, to test agents’ adaptability under reduced reward certainty. Partner-aware agents rapidly achieve high coordination, reaching ∼70% efficiency post-switch, and significantly outperform both other groups. Partner-unaware agents perform sub-optimally, even compared to random agents. Repeated-measures ANOVA confirmed the main effect of agent type (F_2,198_ = 225.34, p < 0.0001). Post hoc Wilcoxon tests showed that partner-aware agents outperformed both partner-unaware and random agents (p < 0.0001). A Friedman test further supported this effect (p < 0.0001). *Right*: Final coordination efficiency after training. Partner-aware agents achieved significantly higher performance than both partner- unaware and random agents (p < 0.0001). Bars represent mean ± SEM; statistical comparisons are indicated above. C. Schematic showing synchronous and asynchronous trial type assignments and quantificationof head bearing angles towards partner rats during asynchronous trials. D. Example trials showing follower and leader gaze dynamics for a WT (*top*) and *Fmr1* (*bottom*) rat. Shaded green area represents when head bearing angle falls below threshold of 40 degrees. E. Within-trial dynamics of gaze variables. A mixed-effects ANOVA revealed significant effects of quartile for follower-only gaze (F_3, 54_ = 11.77, p < 0.001, 𝜂𝑝^2^ = 0.40), but not for mutual gaze (F_3, 54_ = 0.56, p > 0.05 ). Post-hoc comparisons (Holm-corrected) showed that follower- only gaze duration increased from quartile 1 through quartiles 2, 3 and 4 (p < 0.05). No significant effects of genotype or interactions were observed. F. Comparison of proportions of match events for WT and *Fmr1* rats that occurred in asynchronous trials (*left*); session wise comparison of proportion of match events occurring in asynchronous trials (*right*) (*100%:* Mixed-effects linear model, genotype: p > 0.05, session: p < 0.001, genotype X session: p > 0.05. *50%:* Mixed-effects linear model, genotype: p < 0.001, session: p > 0.05, genotype X session: p > 0.05). *** p < 0.001. G. Correlation between mean trial duration and performance for each session. Paired t-test. *** p < 0.001. H. Correlation between normalized follower-only gaze duration and performance for each session. Paired t-test. ** p < 0.01. I. Generalized linear mixed-effects model (GLMM) for assessing effects of gaze predictors on successful coordination (see **Methods** for details) J. Post-hoc comparison (Tukey-adjusted) of effects of different gaze variables during different phases of training on change in log-odds of matching for WT (black) and *Fmr1* (red) rats for 100% reward condition. ** p < 0.01, *** p < 0.001.. K. Same as I, but for 50% reward condition. * p < 0.05, *** p < 0.001.

Building on these model results, we next sought to identify and quantify behavioral markers of peer-directed visual attention in rats performing the cooperative task. To this end, we categorized behavior into two trial types: *synchronous* trials, in which both rats moved in tandem from one well to another (example in **Movie 4**), and *asynchronous* trials (example in **Movie 5**), in which rats initially occupied different reward wells at the beginning of the trial. The asynchronous configuration permitted, though did not necessitate, visual monitoring of the partner, making it particularly suitable for assessing peer-directed attention. We therefore focused on these trials. We began by examining the distribution of cooperative matches across trial types. In WT pairs, matches were more evenly distributed between synchronous and asynchronous trials, whereas *Fmr1* pairs showed a higher proportion of matches occurring during asynchronous trials, particularly under the 50% reward condition (**Fig. 6F**). This indicates that WT matches occurred across both trial contexts, while *Fmr1* matches were more concentrated in asynchronous trials. Examining how asynchronous vs. synchronous match proportions evolved over training, we found that under the 100% reward condition, both genotypes showed a steady increase in match proportions of asynchronous trials across sessions. In contrast, under the 50% reward condition, a clear genotype-specific difference emerged, with *Fmr1* pairs showing a distinct pattern, with higher usage on asynchronous trails for match events (**Fig. 6F**). Next, we found that session-level mean trial duration was negatively correlated with cooperative performance, indicating that shorter, more efficient trials were associated with higher performance (**Fig. 6G**).

We then focused on specific social gaze variables and their contribution to cooperative decisions. We quantified head orientation to estimate for gaze behavior. Head orientation vectors were derived from head-to-neck tracking, and a 40° angular threshold relative to the line of sight was applied to determine whether the partner was within the rat’s binocular field of view (Wallace *et al*., 2013) (**Fig. 6C**, examples in **Movies 5, 6**). Two gaze types were quantified for each trial: follower-only gaze (follower oriented toward leader) and mutual gaze (both rats simultaneously oriented toward each other). Durations of these gaze events were normalized relative to total trial duration. We began by characterizing when rats directed their gaze toward their partners during the task. Across genotypes, partner-directed gaze occurred most frequently in the final quartile of a trial, just prior to departure (**Fig. 6D, 6E**), indicating that visual monitoring was closely tied to movement initiation. Interestingly, follower-only gaze exhibited a weak but significant negative correlation with session performance, suggesting that reliance on follower-only attention decreased in sessions with higher cooperative success (**Fig. 6H**).

Finally, to formally test how different forms of social gaze contributed to cooperative outcomes, we fit mixed-effects generalized linear models (GLMMs) with genotype, training block, and normalized gaze durations as predictors of trial-level cooperation, analyzed separately for asynchronous trials under 100% and 50% reward contingencies (**Fig. 6I**). In our coding scheme, *follower-only gaze* indicates trials in which the follower oriented toward the leader (but not vice versa), whereas *mutual gaze* indicates reciprocal orientation by both rats. Under the 100% reward condition, the models revealed that follower-only gaze became an increasingly strong predictor of successful cooperation across training (**Fig. 6J, Table S1**). Post-hoc comparisons indicated that this effect was driven primarily by *Fmr1* pairs. They continued to rely heavily on the follower attending to the leader to coordinate successfully. In contrast, WT pairs showed a marked reduction in reliance on follower gaze by the final training block, suggesting that coordination became more efficient and less dependent on explicit visual monitoring. A similar pattern was observed for mutual gaze, where the initial reliance diminished over training, consistent with WT pairs transitioning from gaze-dependent to more automatic strategies.

Under the 50% reward condition (**Fig. 6K, Table S2)**, both follower-only and mutual gaze significantly increased the likelihood of coordination overall. However, their use was modulated by genotype and training stage. WT and *Fmr1* pairs both relied on follower-only gaze early in training, but *Fmr1* pairs remained significantly more dependent on the follower monitoring the leader throughout, whereas WT pairs gradually reduced their reliance. For mutual gaze, both genotypes initially benefited from reciprocal monitoring, but this effect declined in later training blocks, again consistent with a shift toward gaze-independent coordination strategies once task contingencies were learned.

Together, these results demonstrate that social gaze in rats, like primates, plays a central role in shaping cooperative outcomes, but that its contribution diverges sharply across genotypes and with training progression. Computational modeling established that partner-aware strategies confer a distinct advantage in cooperative foraging, underscoring the adaptive value of social attention. Behavioral analyses confirmed that rats engaged in peer-directed gaze during asynchronous trials. However, while *Fmr1* rats remained consistently reliant on gaze-dependent strategies across training, WT rats exhibited a dynamic but adaptive shift. They initially deployed gaze as a tool for scaffolding coordination, but progressively transitioned toward a gaze-independent, temporally synchronized strategy. This flexible reduction in gaze reliance highlights an adaptive reorganization of coordination strategies in WT rats, a process that does not emerge in *Fmr1* rats.

### Partner cooperation history shapes cooperative behavior

Social decision-making requires monitoring and adapting to others’ recent actions (Haroush and Williams, 2015; Báez-Mendoza *et al*., 2021). To test whether rats use partner behavior to guide cooperation, and whether this is disrupted in *Fmr1* rats, we analyzed trial-by-trial choices across the two reward conditions (**Fig. 7A**). For the 100% reward contingency, both WT and *Fmr1* rats were more likely to cooperate after their partner had just done so (**Fig. 7B**). In contrast, for the 50% reward contingency only WT rats adjusted their behavior based on recent partner choices.

**Figure 7.**
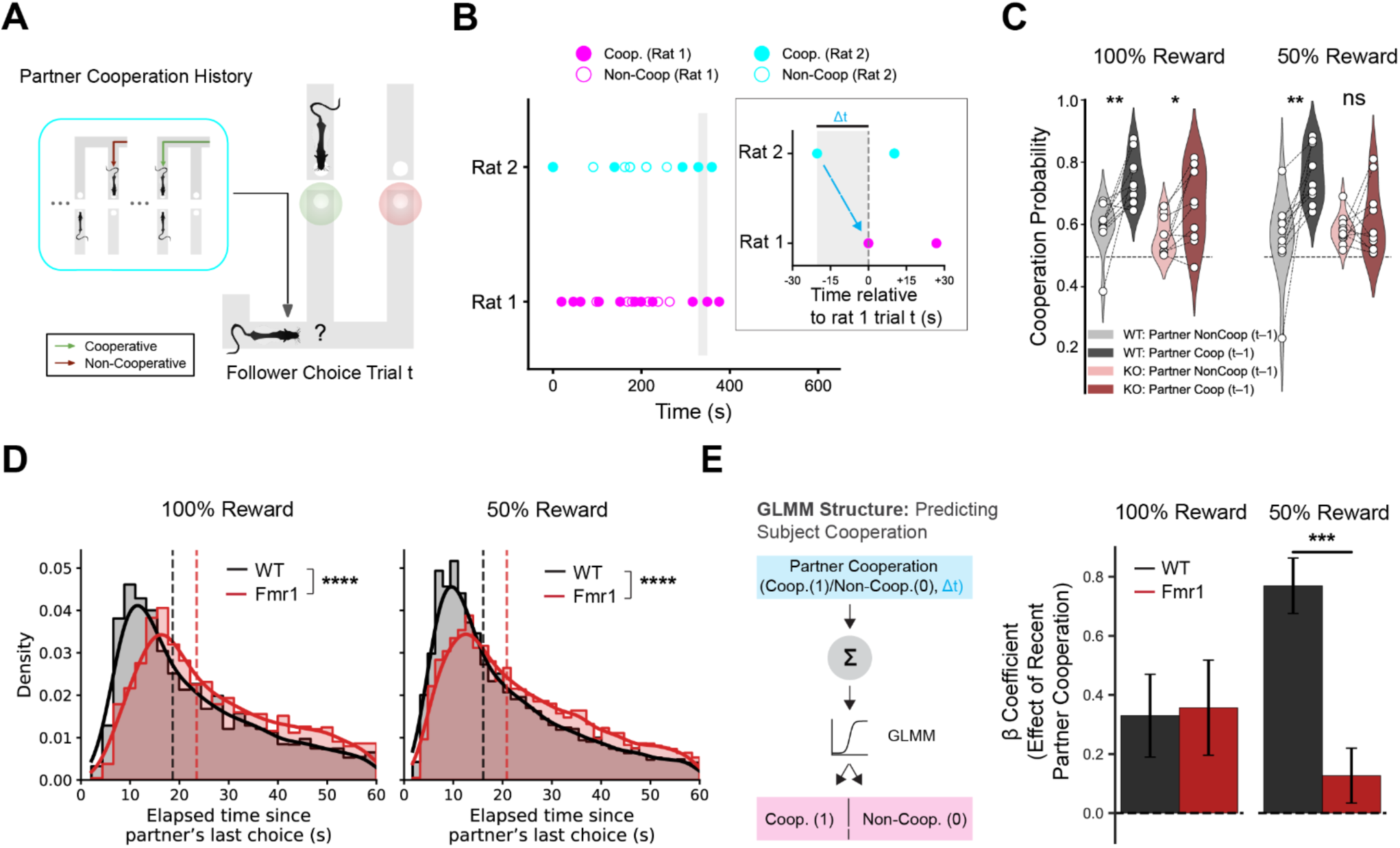
Recent partner choices influence cooperation in WT but not Fmr1^-/y^ rats. A. Schematic of a follower trial. On each trial, one rat (leader) transitions between the arms first; then the other (follower) decides whether to cooperate. B. Timeline of cooperative (filled) and non-cooperative (open) follower choices made by Rat 1 (magenta) and Rat 2 (cyan) in a representative session from a single pair. *Inset:* Zoomed view of a follower trial from Rat 1, showing the most recent cooperative follower choice by Rat 2. Δt denotes the time lag between the rat’s choices. C. Cooperation probability as a function of the partner’s prior choice in WT and *Fmr1*^-/y^ rats under 100% and 50% reward contingencies. Wilcoxon signed-rank test: WT rats, 100%: p < 0.01; *50%:* p < 0.01. Fmr1^-/y^ rats, *100%:* p < 0.05; 50%: p > 0.05. ** p < 0.01, * p < 0.05. *ns*: not significant. D. Distributions show the elapsed time (Δt) between the partner’s most recent choice and the subject’s current trial under the 100% (left) and 50% (right) reward contingencies. Curves are truncated at 60 seconds to highlight behaviorally relevant timescales. Dashed lines indicate genotype-specific medians. Across both contingencies, *Fmr1*^-/y^ rats showed significantly longer partner-referenced Δt intervals than WT rats (100%: U = 6,515,700, p < 0.0001; 50%: U = 4.25×106, p < 0.0001). These timing differences justify the inclusion of Δt and its interactions as covariates in the GLMM to account for baseline differences in decision timing between genotypes. E. GLMM estimating rats’ cooperation based on partner’s previous choice, Δt, and genotype. WT rats show significantly higher contribution of recent partner cooperation in the 50% condition. Fixed-effect β coefficients (±95% CI): WT rats, *100%:* β = 0.36, p < 0.001; *50%:* β = 0.77, p < 0.001. *Fmr1*^-/y^ rats, *100%:* β = 0.26, p < 0.01; *50%:* β = 0.13, p < 0.01. Genotype × partner cooperation interaction at 50%: β = 0.64, p < 0.001. *p < 0.001, p < 0.01, p < 0.05.

To formally assess how partner behavior influenced cooperation, we fit a generalized linear mixed-effects model (GLMM) predicting the likelihood of cooperation on each trial based on the partner’s previous choice, the elapsed time since that choice (Δt), and their interaction (**Fig. 7C**). Including Δt accounted for genotype-related differences in trial timing, such as longer inter-choice intervals in *Fmr1* rats, allowing us to isolate the specific contribution of partner behavior (**Fig. 7D**). To control for individual tendencies, we also included predictors reflecting each rat’s own recent choices. Genotype was included as an interaction term, and random intercepts for rat and session accounted for repeated measures.

The GLMM model confirmed that both WT and *Fmr1* rats were more likely to cooperate following a cooperative partner choice under the 100% reward contingency. However, under the 50% contingency only WT rats adjusted their choices based on recent partner behavior, while *Fmr1* rats did not (**Fig. 7E**).

Together, these findings show that WT rats adapt their cooperative behavior based on recent partner actions, particularly under conditions that require flexible coordination. In contrast, *Fmr1* rats do not, revealing a selective impairment in using social cues to support coordinated decision-making.

## DISCUSSION

In this study, we developed a novel rat spatial cooperative behavior paradigm to investigate social coordination strategies that enable successful cooperation. Our approach provides a simple model, based on the ethologically relevant framework of spatial foraging, for continuous social decision-making in rat dyads under controlled settings, including in a rat model of Fragile X Syndrome. This paradigm allowed us to characterize the coordination-based behavioral organization underlying cooperative behavior and to examine how these processes are disrupted in *Fmr1* rat pairs. We found that both WT and *Fmr1* rat pairs were able to coordinate cooperative visits to the matching reward wells for reward; however, WT rats were more efficient by employing flexible strategies that utilized knowledge of partner choice patterns. *Fmr1* rats relied heavily on a sequential, reactive strategy that involved monitoring partner choices as followers and then executing actions to match partner location, a slower approach that requires asynchronous choices by rats in a pair. In contrast, WT pairs also utilized knowledge of the task structure by coordinating choice sequence patterns with their partner, a predictive strategy that enabled faster, synchronous choices by rats in a pair especially under the probabilistic reward condition. Thus, WT rats are able to employ a complex higher order social coordination strategy for efficient cooperation, which is deficient in *Fmr1* rats.

Our task differs from previous rodent cooperative foraging paradigms (Avital, Aga-Mizrachi and Zubedat, 2016; Jiang *et al*., 2021; Zhang *et al*., 2023; Cheng *et al*., 2025; Erlich *et al*., 2025) in several important aspects. First, since both rats in a pair occupied separate mazes, unlike in more traditional paradigms where animals share the same environment, our design offered several advantages for studying cooperative behavior. By eliminating direct physical interference, the task ensured that coordination arose from genuine monitoring of the partner’s actions rather than from avoidance or displacement. Spatial separation also minimized confounding olfactory cues, which rats could otherwise use to guide behavior. Moreover, having distinct mazes enables precise parametric manipulations of task conditions for each rat, such as reward probability or timing, facilitating a clearer assessment of how one animal’s behavior influences the other. Collectively, these design features strengthen the interpretation that observed coordination reflects strategic, predictive, and socially mediated decision-making rather than simple responses to shared environmental cues. Use of separate mazes will also enable investigation of unambiguous neural correlates of partner locations and actions. Second, by incorporating a 50% reward contingency, we were able to probe how rats flexibly adapted their strategies under conditions where correct choices were only probabilistically rewarded. Finally, reward locations were dynamic rather than fixed, preventing animals from relying on extraneous, task-unrelated cues and instead requiring them to continuously update their behavior in response to their partner’s behavior in real time.

We found that while both WT and *Fmr1* rat pairs learnt to coordinate above chance levels, WT rat pairs outperformed their *Fmr1* counterparts at both 100% and 50% reward contingencies. These findings highlight that, although both WT and *Fmr1* rat pairs were capable of learning to coordinate their choices above chance levels, the efficiency and robustness of cooperative behavior diverged across genotypes. WT pairs consistently outperformed *Fmr1* pairs under both deterministic (100%) and probabilistic (50%) reward contingencies, suggesting that the capacity to flexibly adapt coordination strategies is compromised in *Fmr1* rats. This aligns with evidence that Fragile X Syndrome is associated with impairments in executive function and social adaptability (Budimirovic *et al*., 2006; Klaiman *et al*., 2014; Hooper *et al*., 2018; Schmitt *et al*., 2019; Cregenzán-Royo, Brun- Gasca and Fornieles-Deu, 2022), which may hinder the ability to optimize coordinate. The persistence of coordination in *Fmr1* pairs above chance, however, indicates that basic social engagement mechanisms remain intact, but are insufficient to achieve the same level of efficiency as WT pairs. Together, these results underscore the translational value of our cooperative foraging paradigm for uncovering subtle, context-dependent social decision-making deficits relevant to neurodevelopmental disorders.

Further to ensure that the observed differences in coordination efficiency between WT and *Fmr1* rats were not confounded by basic sensory or associative deficits, we tested both genotypes on a visual cue-association task. Performance accuracy was comparable, indicating that both groups were able to perceive visual stimuli and associate them with reward locations. However, *Fmr1* rats exhibited slower response latencies relative to WT rats reminiscent of altered sensory integration. Importantly, this relative slowness in the visual cue task parallels the broader organization of cooperative foraging behavior in *Fmr1* rats, which unfolded at longer temporal scales compared to the more rapid and synchronous coordination seen in WT pairs. Thus, the cooperative impairments we observed are unlikely to reflect basic sensory or associative limitations, but rather higher-order deficits in executive function and the temporal dynamics of decision-making.

A shift from simultaneous to sequential mode of decision-making, where delaying one’s own actions facilitates incorporating information about a partner’s choice, has been shown to promote coordination (Rapoport, 1997; Brocas, Carrillo and Sachdeva, 2018; Moeller *et al*., 2023). In our task, rats initially relied on such sequential strategies, exhibiting non-zero arrival and departure lags at reward wells relative to their partners. These temporal offsets provided a scaffold for establishing cooperative foraging. As training progressed, WT rats not only became faster overall in their transitions but also developed a flexible mixture of sequential and simultaneous strategies, reflected in reduced lags and the emergence of near-synchronous transitions in cross-correlation analyses.

By contrast, *Fmr1* rats failed to show comparable speeding or flexibility, maintaining elevated lags and lacking short-lag synchrony, suggesting a persistent reliance on sequential strategies without the adaptive shift toward simultaneous coordination that supports efficient cooperation. The use of diverse strategies by WT rat dyads aligns with similar recent reports in a marmoset cooperative lever-pressing task (Meisner *et al*., 2025). Interestingly, recent studies in mice dyads have also shown the use a reactive sequential strategy for cooperative behavior (Cheng *et al*., 2025; Jiang *et al*., 2025). It is possible that the flexible, predictive coordination strategy that we describe here in rat dyads is a hallmark of advanced socio-cognitive abilities.

Furthermore, we observed a dynamic emergence of leader–follower patterns, which arise naturally from the demands of coordination and are particularly prominent in species where social and ecological pressures favor acting together (Krause and Ruxton, 2002). The dynamic leader–follower interactions observed in our rat pairs may reflect a form of spontaneous turn-taking, similar to patterns reported in humans (Clavien and Chapuisat, 2016; Moeller *et al*., 2023). In human dyads, alternating leadership roles facilitates coordination by allowing individuals to anticipate their partner’s actions, maintain joint performance, and avoid conflicts over decisions (Moeller *et al*., 2023). In our task, rats could potentially use a similar strategy as the reward structure did not confer any advantage to the leader in terms of reward acquisition. By flexibly switching which animal leads transitions, they may balance control over cooperative foraging and reduce the likelihood of interference, thereby maintaining efficient coordination over time. While both WT and *Fmr1* pairs exhibited these dynamic patterns, the benefits of such turn-taking may become particularly apparent in contexts requiring rapid, continuous adjustments, highlighting a possible functional advantage of dynamically structured roles in interdependent decision-making.

Our no-vision (NV) control experiments, in which rat pairs were required to coordinate without visual access to one another, underscored the essential role of social vision in sustaining coordination across the task. This finding was reinforced by computational modeling: partner-aware agents, those with access to information about their partner in addition to self- and environmental cues, achieved markedly higher coordination efficiency than agents operating under partner-unaware random policies. Complementary analyses of head direction–based gaze measures further revealed that gaze was especially important for establishing coordination during the early and middle stages of training. By the late training blocks when performance was optimized, WT rats shifted from a primarily gaze-dependent strategy to a more gaze-independent mode, consistent with the emergence of abstract or predictive coordination strategies. The gaze-independent mode depends on maintaining an internal model of current partner strategy and timing, resulting in higher efficiency than a continuous monitoring strategy, and is in marked alignment with similar reports in a marmoset cooperation task (Meisner *et al*., 2025). In this framework, animals in dyads can continue to operate in a coordinated choice pattern regime without requiring continuous monitoring of partner actions, which will only be required to confirm or update the choice patterns and timing in the event of occasional failure to coordinate appropriately. Accordingly, trial duration and follower gaze duration was negatively correlated with performance (**Fig. 6G, H**). In contrast, *Fmr1* rats continued to rely heavily on gaze, indicating a reduced capacity to transition from reactive, visually driven strategies to higher-level predictive control. This is also apparent from the analysis (**Fig. 7**) that shows that partner cooperation history is a significant predictor of ongoing coordination only in WT rats but not in *Fmr1* rats. This persistent reliance highlights a broader deficit in cognitive flexibility and abstraction that may limit their adaptability in social contexts.

Consistent with this interpretation, WT rats displayed higher-order spatial and temporal coordination strategies as training progressed. Their choice sequences became increasingly structured (lower entropy), with a marked rise in the frequency of optimal triplets and the emergence of significant peaks in cross-correlation analyses near zero lag indicating near-synchronous transitions, all of which paralleled their reduced reliance on gaze. *Fmr1* rats developed these strategies to a lesser extent, remaining reliant on gaze-based coordination throughout the task. Together, these findings suggest that WT rats initially employed a reactive strategy grounded in gaze cues, but gradually replaced it with more efficient predictive strategies that involved building internal models of their partner’s behavior. In contrast, the failure of *Fmr1* rats to make this shift points to a broader difficulty in transitioning from externally monitored, effortful strategies to automatic, internalized routines of social coordination.

Overall, our cooperative behavior task provides a tractable model for assessing social interactions and coordination in rodent models. Key features of the paradigm are the relative ease of implementation, reliance on an ethological spatial foraging strategy, detailed quantification of social parameters and coordination strategies, and specificity of deficits observed in *Fmr1* rat models. Further, neurophysiological monitoring and neural circuit manipulation techniques can be utilized in one of both rats in a dyad while navigating two different environments. The task thus provides a foundation to investigate the neural basis of social representations, reciprocal interactions, strategies underlying complex social behavior, and how impairments in these processes can affect social behavior in animal models of autism spectrum disorders.

## ACKNOWLWDGEMENTS

We thank Jadhav lab members for helpful advice and discussions. We also thank Rafael Gabriel for technical assistance with hardware for the visual-cue association task, Audrey Jordan for data collection on the virtual foraging task, and Audrey Hooker for assistance in piloting the behavior task. This work was supported and funded by a SFARI Autism Rat Models Consortium Grant, # SFA-AN-AR-Rat Models-00012700.

## AUTHOR CONTRIBUTIONS

Conceptualization: JHB, SPJ; Experimental Design: AS, ELR, JHB, SPJ; Data Collection: AS, ELR; Data Analysis: AS, ELR; Supervision: SPJ; Writing: AS, ELR, SPJ.

## SUPPLEMENTAL INFORMATION

Figures S1-S2, Tables S1 and S2, and Movies 1-6.

## RESOURCES AVAILABILITY

### Materials Availability

This study did not generate any new unique reagents.

### Data and Code Availability

Data underlying these results will be uploaded in the NWB (Neurodata Without Borders) format to DANDI (ID#TBD) upon acceptance. Code to replicate these results will be available on our lab GitHub (TBD) upon acceptance.

## METHOD DETAILS

### Experimental Model and Subject Details Subjects

Male LE-*Fmr1^em2Mcwi^* rats (*Rattus norvegicus*), hereafter referred to as *Fmr1* rats, and their WT littermates were used in this study. A total of 18 WT and 19 *Fmr1* rats were included, with all subjects aged 3-5 months at the start of behavioral training. Rats were obtained from institutional breeding colonies and were group housed until they were ready for experiments. Prior to starting the experiments and afterwards, rats were single housed under a 12-hour light/dark cycle in a climate- controlled room (temperature: 20°-25°C; humidity: 40% -70%). Food and water were provided *ad libitum*, except during behavioral training, when rats were maintained at approximately 85-90% of their free-feeding weight to ensure their motivation in the task.

Sample size was determined based on [power analysis/preliminary data/statistical justification], ensuring sufficient power to detect biologically meaningful differences in cooperative behavior.

### Ethical Considerations

All experimental procedures were approved by the Institutional Animal Care and Use Committee (IACUC) at Brandeis University (#21001, #24001-A) and conformed to the National Institutes of Health (NIH) guidelines for the care and use of laboratory animals.

## Methods

### Behavioral Apparatus

Rats were trained on elevated W-shaped mazes (80 × 80 cm) with ∼7 cm wide track sections, each consisting of three arms radiating from a central junction. A reward well was positioned at the end of each arm, equipped with an infrared (IR) beam-break sensor for nose-poke detection and automated milk reward delivery. Two mazes were placed in parallel and separated by a transparent acrylic divider, allowing visual and olfactory access while minimizing physical contact between the animals. An overhead color CCD camera (30 fps) was used to record sessions to monitor positions of animals on the mazes.

During behavioral shaping, rats were initially trained on elevated linear tracks (∼80 cm long) equipped with the same IR sensors and reward delivery system to establish nose-poke behavior and familiarize animals with automated reward collection prior to cooperative task training. Behavioral apparatus was controlled using a SpikeGadgets data acquisition system (SpikeGadgets LLC).

### Behavioral Task Design

#### Pre-training and Behavioral Shaping

Prior to training on the cooperative learning task, rats underwent a structured pre-training regimen on linear tracks (80 cm X 7 cm), designed to familiarize them with reward-seeking behavior via nose- poking in designated reward wells. Initially, each rat was trained individually to spatially alternate between the two ends of the track to obtain rewards. This phase encouraged the development of goal-directed navigation and spatial alternation, fundamental skills necessary for subsequent cooperative tasks.

In addition to spatial alternation, rats were trained to execute and maintain nose-pokes within the reward wells with a stepwise increasing hold duration, ranging from 0.2 seconds up to 2 seconds. This incremental hold requirement ensured that the animals learned to sustain their response, a critical component for successful reward acquisition in the cooperative learning paradigm where timing and persistence are essential.

Upon demonstrating proficiency in spatial alternation and poke holding on the linear track, rats progressed to the social phase of behavioral shaping. In this phase, pairs of rats were placed on parallel linear tracks separated by a transparent, perforated acrylic barrier allowing visual, auditory, and olfactory communication while preventing physical contact. Both rats were required to navigate to the same end of their respective tracks and perform simultaneous nose-pokes in the reward wells to obtain rewards. This phase was designed to promote coordinated action and social interaction, facilitating the development of cooperative behaviors critical for the main cooperative learning task.

This gradual shaping protocol ensured that rats were not only trained in individual task components but also conditioned to synchronize their behavior with a partner, thereby providing a robust foundation for assessing cooperative learning in the primary experimental paradigm.

### Cooperative Learning Task

Rat pairs were trained to perform a cooperative learning task that required spatial coordination to obtain joint rewards. Each rat navigated its own W-shaped maze positioned in parallel to its partner’s and separated by a transparent acrylic divider that allowed visual and olfactory access with minimal physical contact. Rewards were delivered only when both rats simultaneously nose-poked at spatially corresponding wells on their respective mazes. The task was continuous and self-paced, enabling extended, naturalistic interactions and the gradual emergence of cooperative strategies over time.

Training began with a 100% reward contingency phase, in which every coordinated match was rewarded. Once rats consistently engaged in the task and demonstrated reliable coordination, we transitioned them to a 50% contingency phase, where coordinated matches were rewarded probabilistically (p = 0.5). This two-phase structure was designed to first reinforce coordination through consistent reward delivery, then challenge animals to sustain cooperative behavior under reduced reward certainty—encouraging behavioral flexibility, persistence, and sensitivity to partner behavior.

Each rat pair underwent at least two training sessions per day, each lasting 20–30 minutes. To minimize potential maze-related biases, rats alternated between mazes across sessions, with assignments pseudorandomized across days.

### Opaque Control Sessions

Following the establishment of proficiency in the coordination task at the 50% reward contingency, rat pairs underwent a series of control sessions designed to isolate the role of visual cues in cooperative behavior. In these sessions, the previously used transparent barrier separating the two rats was replaced with an opaque barrier, effectively preventing the animals from seeing each other. These opaque barrier sessions were strategically interspersed among regular 50% reward sessions with full partner visibility to maintain the integrity of the learned task while allowing within-pair comparisons of cooperative performance under different sensory conditions. Additionally, to eliminate the possibility that rats could rely on auditory signals for cooperation, continuous white noise (∼65 dB) was played throughout the opaque sessions. This auditory masking ensured that any cooperative behavior observed was not influenced by non-visual cues, thus providing a rigorous test of the necessity of visual and auditory communication in supporting coordinated foraging between the rat pairs.

### Visual-Cue Association Task

To assess whether *Fmr1* rats’ deficits in the cooperative learning task stem from impaired social processing or alternative sensory and cognitive impairments, we designed a visually-cued reward association task. This control task evaluated each rat’s ability to detect visual stimuli, associate them with specific actions, and adapt to changing reward contingencies—skills fundamental to successful coordination in the social-W task.

Rats were trained in a W-shaped maze (80 cm × 80 cm) with three reward wells. Each well was equipped with a white LED visual cue, placed on the opposite side of a clear partition (where the partner rat would be located in the main social cooperation task). At any given time, only one well was visually cued, requiring rats to navigate and nose-poke the correct well to obtain a milk liquid reward. Upon correct responses, the LED cue remained active at the same well for ∼2 seconds during reward consumption, before randomly shifting to one of the two remaining wells to promote cue-reward association and facilitate learning.

Training began with a 100% reward contingency phase, where every correct response was rewarded, followed by a 50% reward contingency phase, where correct responses were rewarded with probability 0.5. Sessions were self-paced and fully automated, without a predefined trial structure. Each rat completed 1–2 daily sessions lasting 20–30 minutes, with performance assessed across 30 total sessions. Learning criterion was defined as achieving ≥70–80% correct responses for at least three consecutive sessions before transitioning to the 50% reward contingency phase.

As part of task shaping, rats were pre-trained on a linear track to familiarize them with reward well ports and the association between ports and rewards before full task exposure.

### Cooperation with a Virtual Agent

Rats were trained to navigate a W-shaped maze to obtain milk rewards that were dispensed probabilistically according to a computerized control system. This task was designed to mimic the 50% reward contingency commonly used in cooperative learning paradigm. Specifically, the maze consisted of three arms arranged in a W configuration, each serving as a potential reward location.

On each trial, one of the three arms was designated as the rewarding arm by the computer program, with the selection made randomly to ensure unpredictability. Once a rat received a reward at a particular arm, it was required to move to either of the two remaining arms to obtain the next reward. This rule encouraged the animal to alternate between arms rather than repeatedly returning to the same rewarded location.

The computer program acted as a virtual agent by determining the rewarding arm on every trial, effectively leading the sequence of trials. Rats were therefore challenged to learn and anticipate the virtual agent’s random reward assignment to optimize their foraging strategy. This setup required the rats to either guess the rewarding location or cooperate implicitly with the virtual agent’s programmed contingencies, enabling the study of decision-making processes under probabilistic reward conditions

### Behavioral Quantification and Statistical Analysis

Unless noted otherwise, behavioral analyses used the first 21 sessions for the 100% reward contingency and the first 35 sessions for the 50% reward contingency, matching the minimum available sessions across all pairs.

### Performance in Cooperative Task

Behavioral performance was assessed by calculating the ratio of coordinated matches to the total number of arm transitions made by each rat pair. This measure controlled for variability in task engagement across sessions and genotypes. Performance was expressed as the percentage of matches relative to total transitions:

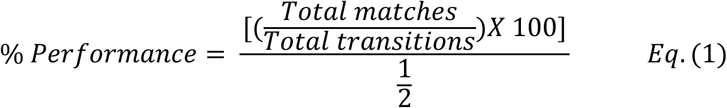

For an optimally performing pair, each coordinated match would require at least two transitions— one from each rat—resulting in a maximum achievable performance score of 100%. Behavioral performance was quantified as percent correct responses per session, with additional analyses of latency to correct choice and performance adaptation between 100% and 50% reward contingencies.

### Performance Quantification and Statistical Analysis for Visual-Cue Association Task

Behavioral performance was quantified as percent correct responses per session, with additional analyses of latency to correct choice and performance adaptation between 100% and 50% reward contingencies. A mixed-effects model was used to assess learning trajectories, incorporating random effects for individual rats and fixed effects for session, genotype, and reward contingency.

Statistical comparisons included:

● Main effect of session (learning progression; p < 0.001),
● Main effect of genotype (no significant differences; p = 0.981, indicating preserved visuo-cognitive performance in *Fmr1* rats),
● Genotype × Session interaction (not significant; p > 0.05, suggesting similar learning trajectories between groups).

Performance data were smoothed using a Gaussian kernel (σ = 2 sessions) for visualization, and multiple comparisons were corrected using Bonferroni/FDR correction.

### Performance in Cooperation with a Virtual Agent

Behavioral performance was quantified as percent of correct rewarded responses per session.

### Position tracking

DeepLabCut 2.3.9 (Mathis *et al*., 2018) was utilized to track the position of different body parts in videos of rats performing the behavioral task. Prior to model training, each video recording was processed to generate two separate datasets, corresponding to the rat located on the left W-maze and the rat located on the right W-maze. This involved cropping the original videos to isolate and extract the respective rat’s movements within each W-maze. 20-40 frames from at least 10 videos (from each experimental cohort) were automatically extracted (using *k-means algorithm*) for labelling the keypoints and training the model, thus capturing the variability in posture and luminance conditions. The extracted frames were manually labeled to define the keypoints corresponding to different body parts. A ResNet-50 based model was trained with 95% of the labelled data with default parameters for 100,000 number of training iterations. Keypoints with likelihood < 0.6 were set to NaN, interpolated, and the resulting positions smoothed with a Gaussian kernel (σ = 150 ms). We then used a p-cutoff of 0.6 to condition the X,Y coordinates for future analysis. This network was then used to analyze videos from similar experimental settings.

### Quantification of Peer-Directed Head Orientation Events

To quantify social attention, we analyzed partner-directed head orientation using keypoint tracking data from DeepLabCut. Analysis was restricted to time periods meeting two criteria: (1) the rats were positioned at distinct reward wells on their respective W-mazes, ensuring spatial separation; and (2) the focal rat was classified as the Follower if it initiated a transition to a new reward well while the partner remained stationary, thereby making a cooperative or non-cooperative choice. A trial was defined as the interval beginning when the rats were in this configuration and ending when the Follower departed its well to initiate a choice. For each trial, we quantified partner-directed head orientation from the Follower to the Leader, from the Leader to the Follower, and mutually.

We calculated a head direction vector (v^head^) for the focal rat, defined from the neck to the head center, and a partner vector (v^partner^), defined from the focal rat’s head center to the partner’s body center.

The bearing angle 𝜃 was computed using:

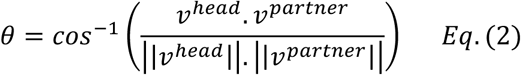

A head orientation event was classified as peer-directed if θ fell below a predefined angular threshold, indicating that the focal rat was directing its head toward the partner. We adopted an angular threshold of 40°, in line with criteria used by (Wallace *et al*., 2013). These supplementary analyses yielded qualitatively similar results, indicating that our findings were not overly sensitive to the specific threshold choice.

We then extracted three key summary metrics: (1) the proportion of time the Follower rat oriented toward the Leader (g_F_) and (2) the proportion of time both rats simultaneously oriented toward each other (g_M_). These metrics quantified the prevalence and mutuality of social attention between the dyad members.

### Model structure

The mixed-effects logistic regression model included:

All main effects for gaze measures, genotype, and session blocks, all two-way and three-way interactions between predictors, random intercepts for individual animals, random intercepts for sessions nested within animals.

The model was specified as:

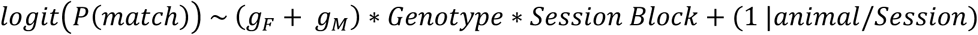

where:

𝑔*_r_*, and 𝑔_M_ denote normalized follower-only and mutual gaze durations, respectively.

𝐺𝑒𝑛𝑜𝑡𝑦𝑝𝑒 denote the genotype of the rat.

𝑆𝑒𝑠𝑠𝑖𝑜𝑛 𝐵𝑙𝑜𝑐𝑘 denotes the session block (early, mid, late) nested within animal.

### Statistical implementation

All analyses were conducted in R (version 4.4.3). The mixed-effects model was fitted using the function from the *glmmTNB* package with the binomial family and logit link function. Model convergence was verified.

### Post-hoc analyses

To interpret significant interactions, we conducted post-hoc contrast analyses using the *emmeans* package. Specifically:

1. **Within-genotype contrasts**: We computed estimated marginal trends (*emtrends*) for each gaze measure across session blocks within each genotype, followed by pairwise contrasts to test for significant changes across training phases.
2. **Between-genotype contrasts**: We compared estimated marginal trends between genotypes within each session block to identify when genotype differences emerged.

All post-hoc comparisons were adjusted for multiple testing using the Tukey method. Statistical significance was set at α = 0.05 for all tests.

### Model evaluation

Model fit was assessed using marginal R² (variance explained by fixed effects only) and conditional R² (variance explained by the complete model including random effects). Results are reported as odds ratios with 95% confidence intervals and p-values.

### Assignment of Leader and Follower Identities for Matched Events

For each matched event, the rat that arrived first at the reward well on its respective W-maze arm was designated as the Leader, while the other rat was assigned as the Follower. These roles were determined based on the precise timing of their reward well entries (poke timings).

### State-Space Modeling of Leadership Probability

To estimate the dynamic probability of a rat leading during each matched event across sessions, we applied a state-space modeling approach to the binary leadership vector derived for each rat. The vector encoded leadership status per event, with 1 indicating the rat led the event and 0 indicating it followed.

Following the framework described by Smith et al. (2009), we modeled the latent, time-varying propensity to lead as a hidden continuous state that evolves according to a Gaussian random walk. Observed binary leadership outcomes were assumed to arise from a Bernoulli process with a logistic link function mapping the latent state to a probability of leading.

Parameter inference was performed using an expectation-maximization (EM) algorithm combined with a Newton-Raphson procedure to update latent states. This algorithm iteratively estimated the smoothed latent state trajectory, the associated posterior variance, and the time-varying probability of leading, while updating process noise parameters until convergence.

Confidence intervals for the latent state and the resulting probability estimates were calculated using numerically integrated posterior distributions. The model was implemented with performance optimizations including Just-In-Time compilation (Numba) and batch processing of confidence limits.

In brief, this approach allowed for the quantification of uncertainty in leadership probability over time, yielding smooth, probabilistic estimates of leadership dynamics across matched events within and between sessions.

We implemented a state-space model following Smith et al. (2009) to estimate the latent, time- varying probability of leadership from binary observations of leadership per matched event.

The hidden state 𝑥_k_ evolves as a Gaussian random walk:

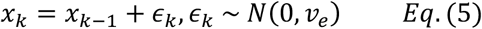

where 𝑣_e_ is the process noise variance.

The observed binary leadership outcome 𝑛_k_∈ {0,1} at time𝑘 is modeled as a Bernoulli trial with probability:

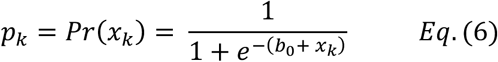

The goal is to estimate the posterior distribution of the latent states {𝑥_𝑘_} and process noise 𝑣_e_ given the observations {𝑛_𝑘_}. Due to the nonlinear logistic observation model, exact inference is intractable.

So, we used an Expectation-Maximization (EM) algorithm combined with a Newton-Raphson update at each time step to iteratively estimate the latent states and optimize 𝑣_e__eve. The Newton-Raphson step solves:

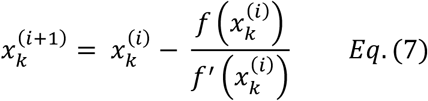

where,

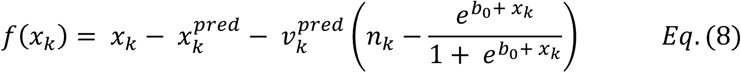

and

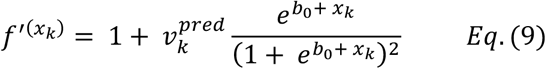

Here, 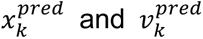 are the predicted state mean and variance from the previous iteration. The model yields smoothed estimates of the latent state 𝑥*_k_*, from which the time-varying leadership probability 𝑝*_k_* and confidence intervals are computed.

### Choice Triplet Counts, Exploration Efficiency, and Sequence Entropy

To investigate patterns of exploration and choice regularity, we analyzed the sequences of reward well visits made by individual rats during the task. Each reward well poke was recorded and used to reconstruct the animal’s choice sequence over time. From these data, we defined *choice triplets* as contiguous sequences of three consecutive well visits, representing the smallest temporal unit that could capture structured exploration across multiple spatial targets.

A sliding non-overlapping window approach was used to extract all such triplets within each behavioral session for every rat. Given the three reward wells (labeled 1, 2, and 3), eight unique choice triplets were possible: 1-2-1, 1-2-3, 1-3-1, 1-3-2, 2-1-2, 2-3-2, 3-1-3, and 3-2-3. Among these, only two triplets, 1-2-3 and 1-3-2, resulted in visits to all three wells within the span of three pokes, thereby representing maximally efficient exploration of the available reward space. These were classified as *optimal choice triplets*. The remaining six patterns, which revisited wells before sampling all three, were designated as *suboptimal triplets* as they reflect redundant or inefficient exploration paths. For each rat and each session, we quantified the frequency of occurrence of each triplet type. This allowed us to characterize how often rats explored all available reward options in a systematic manner versus exhibiting stereotyped or repetitive behaviors.

To directly quantify efficiency of exploration, we assessed how quickly and frequently each rat visited all three reward wells during a session. The rationale is that an optimally exploring rat would systematically sample the environment, rapidly visiting all wells without unnecessary repetition. In the ideal case, such a rat would encounter all three wells after just three visits, for example via triplets like *1-2-3* or *1-3-2*, and would do so repeatedly across the session.

In contrast, a rat with poor exploration efficiency might display repetitive or biased behavior, such as persistently alternating between only two wells (e.g., *1-2-1*), and take more visits to cover the entire reward space. These inefficient patterns are reflected in a higher frequency of suboptimal triplets.

To operate this, we computed the proportion of optimal triplets (i.e., *1-2-3* and *1-3-2*) over the total number of triplets in each session:

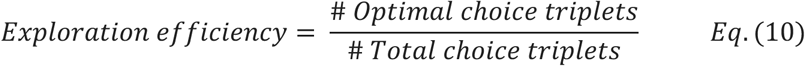

This proportion served as an index of exploratory efficiency. Higher values indicated systematic and balanced foraging across all wells, while lower values reflected redundancy or bias in choice behavior. Session-wise comparison of this metric thus provided a dynamic measure of how well the rats engaged in efficient sampling of the reward space during the coordination task.

In parallel, we also quantified the entropy of choice sequences as a separate measure of behavioral regularity. For each session, we computed the sample entropy of the full choice sequence (i.e., the sequence of visited reward wells) to capture the degree of predictability in the animal’s behavior. A reduction in entropy over sessions would suggest increasingly stereotyped or regularized behavior, while persistently high entropy would reflect continued exploratory variability. Entropy values were computed for each session per rat and subsequently compared across genotypes.

### Cross-Rat Choice Sequence Similarity Analysis

To quantify behavioral choice sequence similarity between rats within a pair, we computed cross-rat Jaccard similarity matrices. For each rat pair, all sessions of the first rat were compared against all sessions of the second rat. Each matrix entry corresponded to the Jaccard similarity between the two corresponding sequences, defined as:

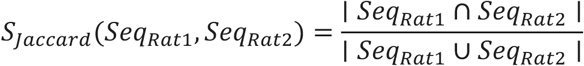

where 𝑆𝑒𝑞_Rat1_and 𝑆𝑒𝑞_Rat2_ are the sequences of maze arm transitions for a given session of each rat. Values range from 0 (no shared transitions) to 1 (identical sequences), providing a direct measure of behavioral alignment through similarity of choice sequences between paired rats.

For visualization, lower triangular versions of cross-rat similarity matrices were displayed as heatmaps, with sessions of one rat along the x-axis and sessions of the partner along the y-axis. A color gradient indicated the magnitude of similarity.

### Multinomial Logistic Regression and Cross-Validation for Quantifying Coordination Between Rats

As another measure of degree of coordination between pairs of rats during the cooperative learning task on W-mazes, we implemented a multinomial logistic regression approach to evaluate how well a rat’s next choice could be predicted from its partner’s current location.

To assess whether a rat’s location could predict its partner’s subsequent choice, we trained multinomial logistic regression classifiers (using *scikit-learn* Python package) to predict the rat’s next choice on its own maze based on the current position of its partner located on the other maze. The input predictor was one-hot encoded, and models were evaluated using 5-fold stratified cross- validation for each rat individually. Classification accuracy was computed and averaged across folds to generate a rat-wise predictive accuracy measure.

To evaluate whether the observed accuracies were greater than expected by chance, we conducted permutation testing by independently shuffling the vectors of the focal rat’s next choices and current locations of the partner rat 1000 times for each rat and repeating the cross-validation procedure. The resulting null distribution of accuracies was used to compute empirical p-values for each group by comparing observed mean accuracy to the distribution of permuted accuracies.

Group-wise differences in classification accuracy (both observed and null) between WT and *Fmr1* rats were evaluated using the Mann–Whitney U test (two-tailed).

### Permutation Testing of Coordination Performance

To test whether coordination improvements reflect true learning, we compared observed performance to null distributions generated by circularly time-shifting one animal’s entire sequence of reward well visits within each session, thereby preserving session-level statistics while disrupting coordinated timing between animals. Each null distribution was constructed from 1,000 permutations. P-values reflect the proportion of permutations in which surrogate performance matched or exceeded the observed value.

### Partner Behavior History Analysis

To test whether rats adjust their cooperative decisions based on recent partner behavior–and whether this differs by genotype–we analyzed trial-by-trial choices during “follower trials”. In each trial, one rat (the leader) enters an arm first; the other rat (the follower) then decides whether to choose the matching arm and cooperate. Leader and follower roles alternate dynamically within each session, allowing both rats to serve in both roles.

For each follower trial, we aligned the partner’s most recent choice (from their previous follower trial) and computed the elapsed time between that choice and the current follower decision (Δt) for use in the statistical model. Trial sequences for each rat were constructed by sorting all trials chronologically and identifying those where the rat served as Follower. Partner history variables were derived by matching each follower trial to the most recent follower trial completed by the partner.

### Generalized Linear Mixed-Effects Modeling of Partner Influence

To quantify how recent partner behavior influenced cooperative decisions—and how this relationship differed by genotype—we fit a generalized linear mixed-effects model (GLMM) predicting the likelihood of cooperation on each follower trial (binary outcome: cooperate = 1, not cooperate = 0).

Fixed effects included the partner’s previous choice (binary), the elapsed time since that choice (Δt, continuous), and their interaction. Genotype was modeled as an interaction term with all predictors to assess genotype-specific effects.

We also included covariates reflecting the subject’s own recent choice history to control for individual behavioral tendencies. Δt was included to account for inter-trial timing differences, particularly the longer partner-to-subject intervals observed in *Fmr1* rats.

Random intercepts for rats and sessions accounted for repeated measures. Separate models were fit for the 100% and 50% reward contingencies to isolate effects within each condition.

Models were implemented in Python using the *pymer4* package (v0.8) and validated against equivalent R-based implementations (*lme4*).

### Multi-Agent Reinforcement Learning Framework for Modeling Cooperative Behavior

To complement behavioral findings and test whether access to partner information facilitates cooperative behavior, we developed a multi-agent reinforcement learning (RL) model of the spatial coordination task (Tan, 1993; Du *et al*., 2023; Tsutsui *et al*., 2024). In this model, two agents independently navigated discrete three-location grids and were required to coordinate their choices to receive a shared reward. We implemented three agent types with varying access to task-relevant cues: **partner-aware agents** received both self- and partner-location information; **partner-unaware agents** received only self-related location information; and **random agents** received no observations and selected actions uniformly at random. This design allowed us to isolate the role of partner information in driving coordination over the course of learning.

The environment consisted of two agents, each occupying its own discrete 1D grid composed of three locations, representing the reward wells in the behavioral task. On each timestep, one agent was randomly selected to act first. This agent observed the current environmental state—including its current and previous locations, the location of the previously rewarded match, and, for partner-aware agents, the partner’s current location—and then selected an action. The second agent subsequently observed the updated environment state, which included the same set of features along with the first agent’s new position, and selected its action accordingly. A joint reward of +1 was delivered only if both agents selected the same location and that location differed from the previously rewarded location. Random agents received no observations and chose actions randomly. This framework preserved the task’s turn-taking dynamics and non-repetition rule while allowing precise manipulation of partner cue access.

Partner-aware and partner-unaware agents learned using standard tabular Q-learning. For each agent, a state–action value table was maintained and updated based on the following rule:

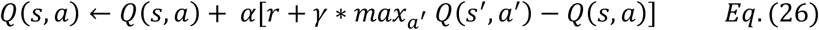

where 𝑄(𝑠, 𝑎) is the estimated value of taking action 𝑎 in state 𝑠, 𝛼 is the learning rate, 𝛾 is the discount factor, 𝑟 is the reward received, and 𝑠, is the resulting state after the action. Agents selected actions using an epsilon-greedy policy to balance exploration and exploitation. The probability of random exploration (epsilon) was decayed exponentially across episodes to encourage early exploration and later convergence on learned policies. Specifically, epsilon decayed from 1.0 to 0.01 over 90% of the training episodes according to:

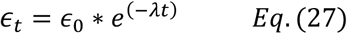

where 𝜖_0_is the initial exploration rate and 𝜆 is the decay rate calculated to ensure that 𝜖_t_= 0.01 by the end of the decay window. After this point, epsilon was held constant at 0.01 for the remainder of training.

### Statistical Analysis

All statistical analyses were conducted using Python (scipy, statsmodels) or RStudio (lme4, afex, effects, emmeans, glmmTNB).

## SUPPLEMENTARY FIGURES

**Figure 1-Supplementary 1:**
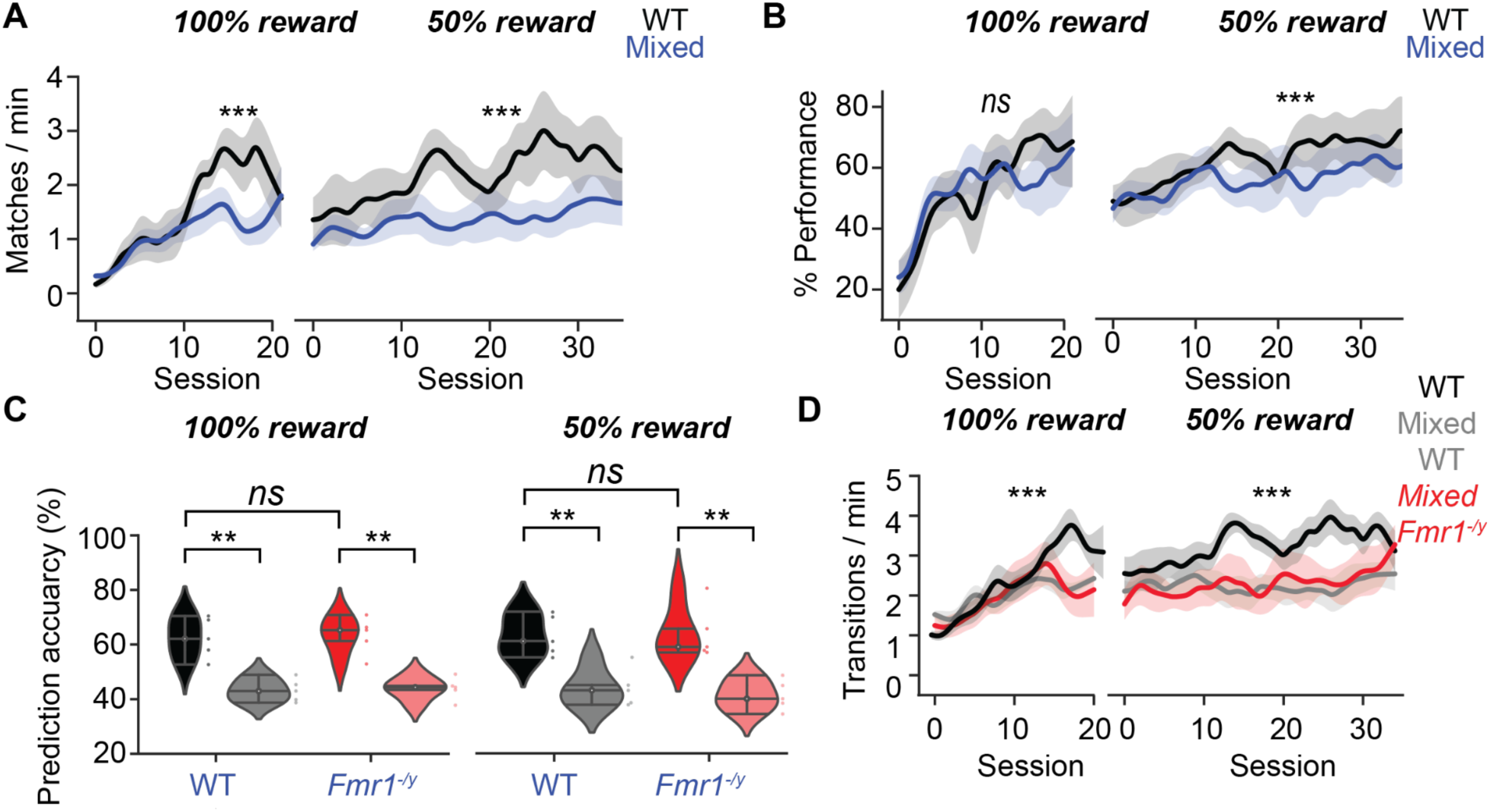
Coordination efficiency and task engagement for mixed genotype pairs. A. Match rates for WT and mixed pairs for 100% (*left*) and 50% (*right*) reward contingencies. *100%:* Two-way RM ANOVA (genotype: p < 0.001, session: p < 0.001, genotype X session: p < 0.05). *50%:* Two-way RM ANOVA (genotype: p < 0.001, session: p > 0.05, genotype X session: p > 0.05). *** p < 0.001. B. Performance metrics for WT and mixed pairs in 100% (*left*) and 50% (*right*) reward contingencies. *100%:* Two-way RM ANOVA (genotype: p > 0.05, session: p < 0.001, genotype X session: p > 0.05). *50%:* Two-way RM ANOVA (genotype: p < 0.05, session: p > 0.05, genotype X session: p > 0.05). *** p < 0.001. C. Prediction accuracy of rats’ next choice given partner’s choice based on a multinomial GLM (5-fold cross-validation) in 100% (*left*) and 50% (*right*) reward contingencies (Mann-Whitney U test, * p < 0.05, *** p < 0.001). Transition rates for WT (pure WT pairs) and WT *Fmr1^-/y^* rats (mixed genotype pairs) under 100% (*left*) and 50% (*right*) reward contingencies. *100%:* Two-way RM ANOVA (genotype: p < 0.001, session: p < 0.001, genotype X session: p > 0.05). *50%:* Two-way RM ANOVA (genotype: p < 0.001, session: p > 0.05, genotype X session: p > 0.05). *** p < 0.001.

**Figure 5-Supplementary 1:**
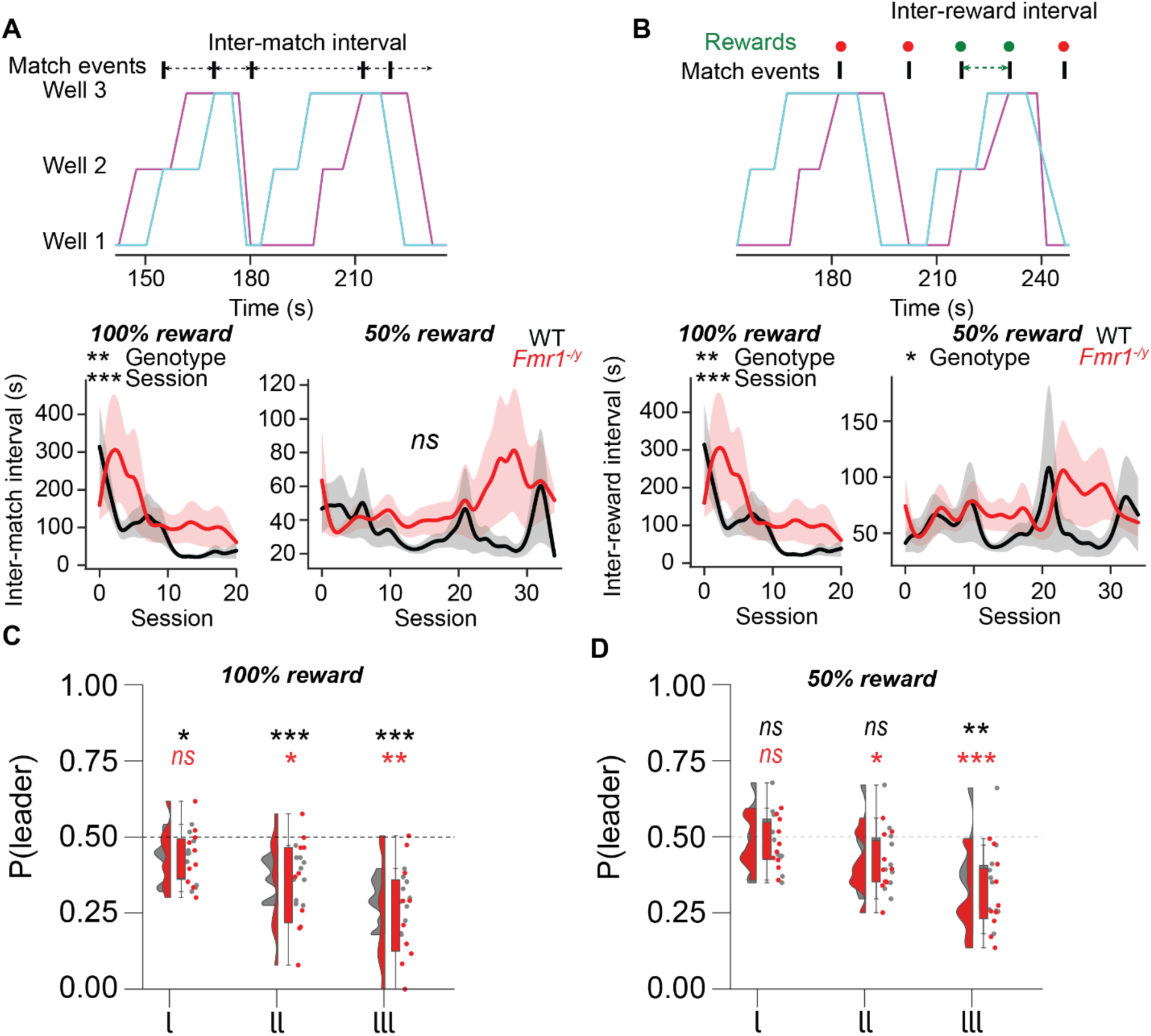
Timescales of behavioral coordination. A. *Top:* Snippet showing intervals between successive cooperative events. *Bottom:* Mean interval between successive cooperative events in WT and *Fmr^1-/y^* pairs for 100% (*left*) and 50% reward contingencies (*right*). *100%:* Two-way RM ANOVA (genotype: p < 0.01, session: p < 0.001, genotype X session: p > 0.05). *50%:* Two-way RM ANOVA (genotype: p > 0.05, session: p > 0.05, genotype X session: p > 0.05). ** p , 0.01, *** p < 0.001. B. *Top:* Snippet showing intervals between successive rewarded cooperative events. *Bottom:* Mean interval successive rewarded cooperative events for WT and *Fmr^1-/y^* rats for 100% (*left*) and 50% reward contingencies (*right*). *100%:* Two-way RM ANOVA (genotype: p < 0.01, session: p < 0.001, genotype X session: p > 0.05). *50%:* Two-way RM ANOVA (genotype: p < 0.05, session: p > 0.05, genotype X session: p > 0.05). * p < 0.05, ** p < 0.01, *** p < 0.001. C. Conditional probabilities of assuming leader role after one, two or three successive leader roles for WT and *Fmr1^-/y^* rats in 100% reward contingency. Wilcoxon test (*ns* p > 0.05, * p >0.05, ** p <0.01, *** p <0.001). D. Conditional probabilities of assuming leader role after one, two or three successive leader roles for WT and *Fmr1^-/y^* rats in 50% reward contingency. Wilcoxon test (*ns* p > 0.05, * p >0.05, ** p <0.01, *** p <0.001).

**Table S1:**
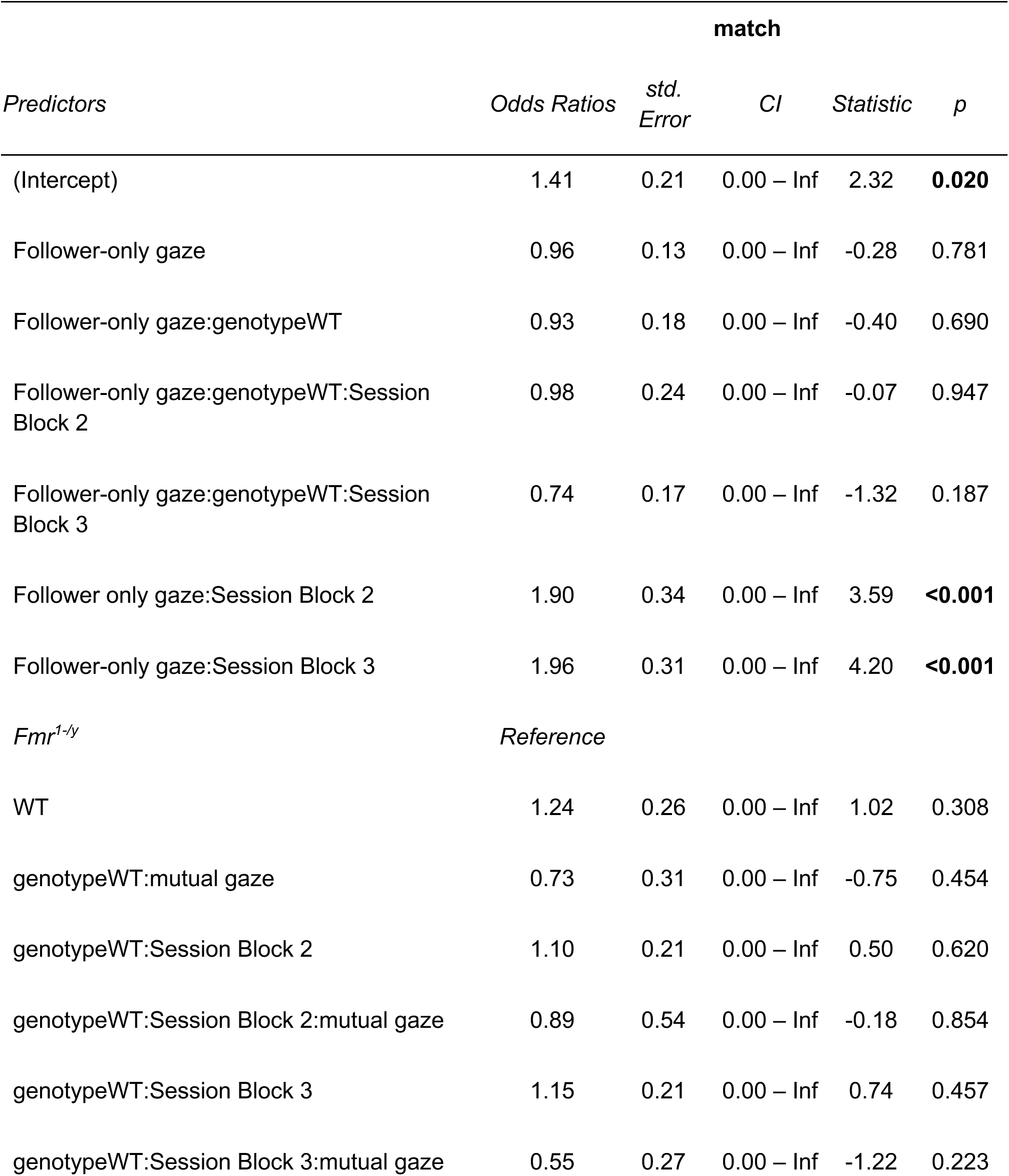
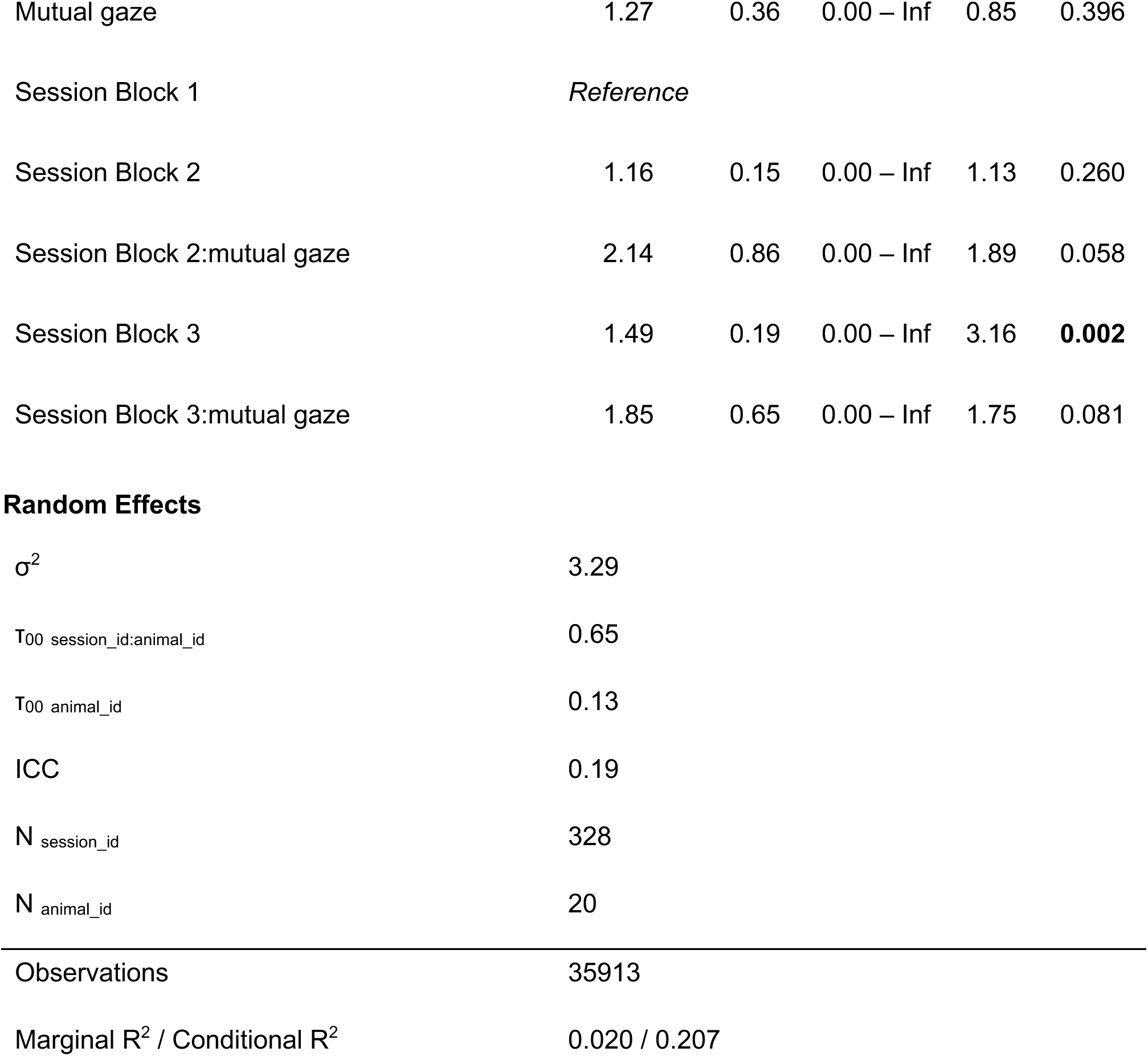
Mixed-Effects Logistic Regression Results (100% reward condition)

**Table S2:**
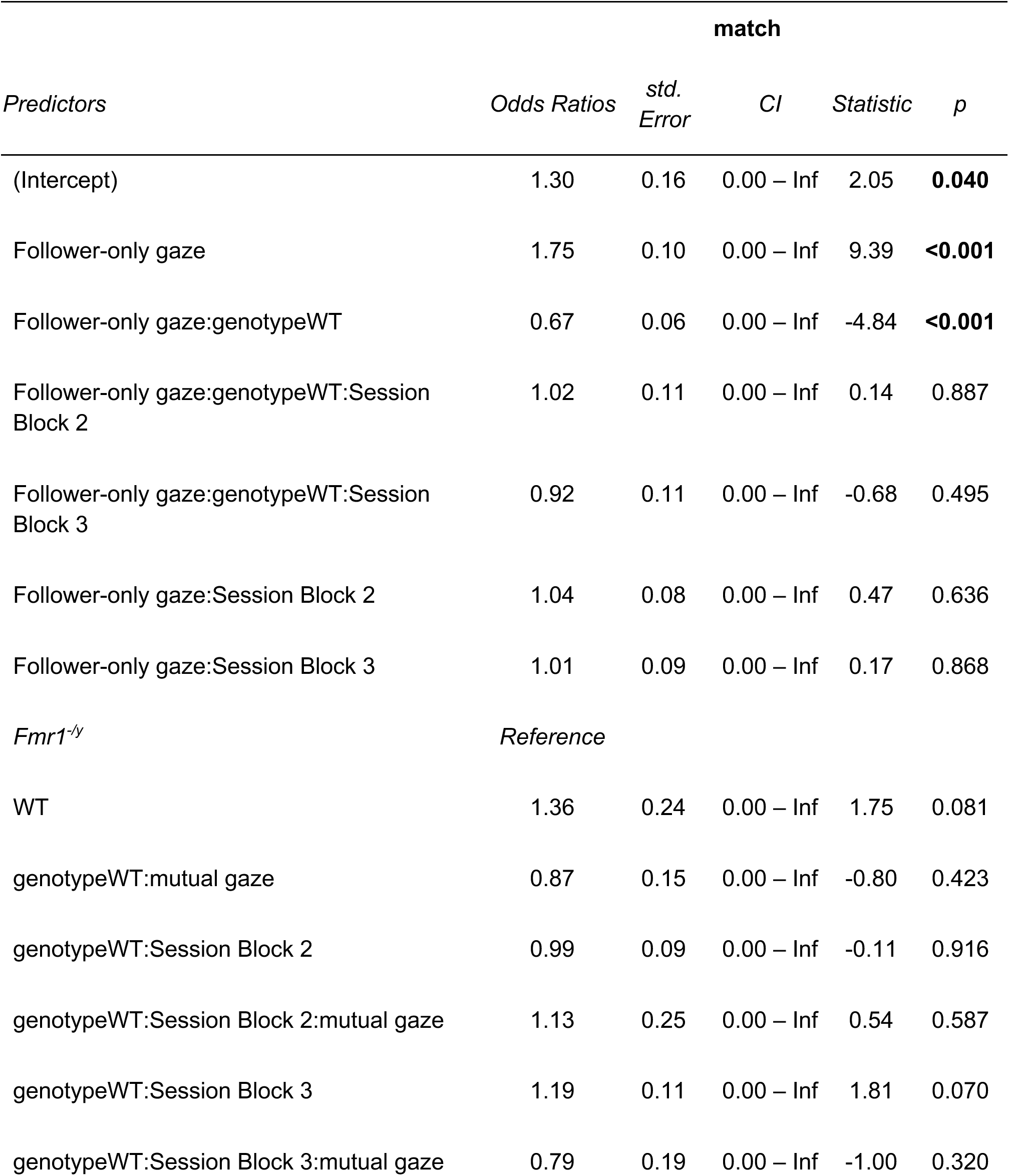
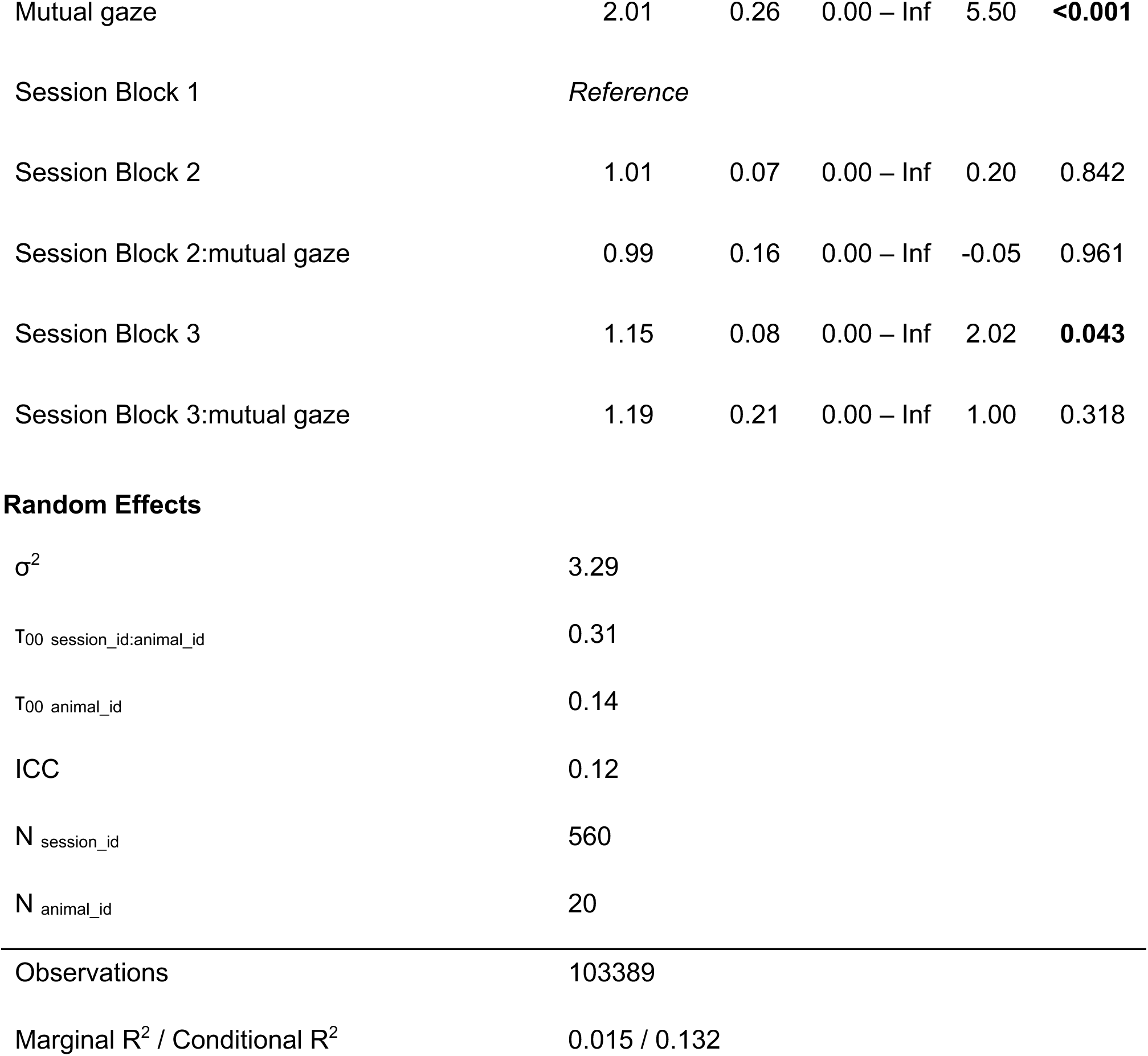
Mixed-Effects Logistic Regression Results (50% reward condition)

